# Cerebral hemodynamics: a mathematical model including autoregulation, baroreflex and extracranial peripheral circulation

**DOI:** 10.1101/2021.06.11.448061

**Authors:** Francisco A. Garcia, Deusdedit L. Spavieri, Andreas Linninger

## Abstract

Cerebral autoregulation, the physiological capability to regulate cerebral blood flow, may be assisted by short-term mean arterial pressure control via baroreflex, which, among several effects, modulates total peripheral resistance. It is unclear, however, whether the resistance of the head and neck vasculatures is also affected by baroreflex and whether these extracranial vessels assist autoregulation. Since sensing technologies such as functional Near-Infrared Spectroscopy and noninvasive intracranial pressure monitoring by strain gauge may be influenced by superficial tissue, it is clinically relevant to understand the relations between intracranial and extracranial hemodynamics. Therefore, we created an autoregulation model consisting of arteries and arterioles regulated by the intralumial pressure and microcirculation regulated by local blood flow. As the first critical step to quantify the signal deterioration introduced by the extracranial circulation on superficial sensors, the extracranial peripheral circulation of the head and neck and baroreflex regulation of the peripheral vasculature and of heart rate were also included. During simulations of a bout of acute hypotension, the model predicts a rapid return of cerebral blood flow to baseline levels and a prolonged suppression of the blood flow to the external carotid vasculature, in accordance with experimental evidence. The inclusion of peripheral control via baroreflex at the external carotid vasculature did not assist cerebral autoregulation, thus we raise the hypothesis that baroreflex may act on the head and neck vasculatures but this action has negligible effects on regulation of cerebral blood flow. When autoregulation is impaired, results suggest that the blood flow of the brain and of the head and neck present similar dynamics, while they are weakly coupled when autoregulation is intact. The model also provides a mechanistic explanation of the protection brought by cerebral autoregulation to the microvasculature and to the brain parenchyma. Our model forms the foundation for predicting the interference introduced by the superficial tissue to nonivasive sensors.

## Introduction

Cerebral autoregulation (CA) is believed to comprise a multitude of cerebrovascular and neurogenic mechanisms that keep cerebral blood flow (CBF) in normal ranges despite variations in the cerebral perfusion pressure from 50 to 150 mmHg^1^. Ample evidence suggests that the systemic blood pressure regulation via baroreflex plays a role in maintaining CBF^2–4^. Baroreflex acts for example on the total peripheral resistance by modulating sympathetic nerve activity. However, very little is known about the sympathetic effect on the vascular bed of the external carotid artery (ECA) and the dynamic relations between the intracranial dynamics and the extracranial circulation of the head and neck^4^. The internal (ICA) and external carotid arteries are physically coupled at the common carotid (CCA) bifurcation. The results of acute hypotension experiments induced by thigh-cuff inflation-deflation initially indicated that the vascular bed of the head and neck constrict due to an increase in sympathetic tone via baroreflex, thus contributing to the reestablishment of CBF^3^. However, more recent work suggests that the vascular bed of the ECA is not regulated by sympathetic activity and does not contribute to CBF regulation during acute hypotension in healthy young men^4^. More background on the CA and baroreflex mechanisms is given in Supplementary Section 1.

Episodes of hypotension^5^ often arise to neurointensive care patients^6^. If their CA is impaired, as frequently happens^7^, ischemic brain damage can occur^8^. To monitor cerebral state of critically ill patients, usually adopted technologies include invasive intracranial pressure monitoring and noninvasive CBF assessment via transcranial doppler. Increasing attention is being given to other noninvasive surface technologies such as functional Near-Infrared Spectroscopy and noninvasive intracranial pressure (ICP) monitoring by strain gauge^9^. Signal acquisition of these sensors, however, might be influenced by the superficial tissue^9–12^. To test this possible interference, a mathematical model that accounts for the most relevant intracranial and extracranial dynamics of the head and neck is a potential tool. The model may also be used to predict the role of the extracranial vasculature of the head and neck to CBF regulation and the effects of dynamic changes of mean arterial pressure (MAP) to the brain, such as during episodes of hypotension that are clinically important in patient care.

Here we present a model of the intracranial dynamics including CA mechanisms that capture the main factors as function of the sizes of the vessels, baroreflex control of peripheral vasculature and of heart rate (HR) and the peripheral circulation of the head and neck, which may act as a compensatory blood reserve to CBF. Building on earlier work^13–16^, the model is able to simulate pulsatile ICP, incorporating the main compartments of the cerebrospinal fluid (CSF) system and the brain parenchyma. The justification of the model lies on the finding that prior detailed models of the CSF system and brain parenchyma dynamics were not intended to include regulatory mechanisms, while models of baroreflex and of CA offer oversimplified descriptions of the CSF and brain parenchyma systems. A comparative review of previous work on CA, baroreflex and intracranial dynamics is given in Supplementary Tables S2 and S3. Although some excellent models have been proposed (e.g.^17–20^), an unifying model with a detailed description of the intracranial dynamics including regulatory mechanisms and the extracranial vasculature of the head and neck has not yet been seen in the literature. Our model includes the extracranial circulation of the head and neck regulated by baroreflex and the intracranial blood, CSF and brain parenchyma systems, derived from the Linninger model^13^. Cerebral autoregulation was modelled as being regulated by the intralumial pressure at the arteries and arterioles, and regulated by local blood flow at the microcirculation. Baroreflex control of the peripheral vasculature and of heart rate were adapted from Lau-Figueroa model^14^. An overview of the model is given in Fig 1 (**A**).

**Figure 1.**
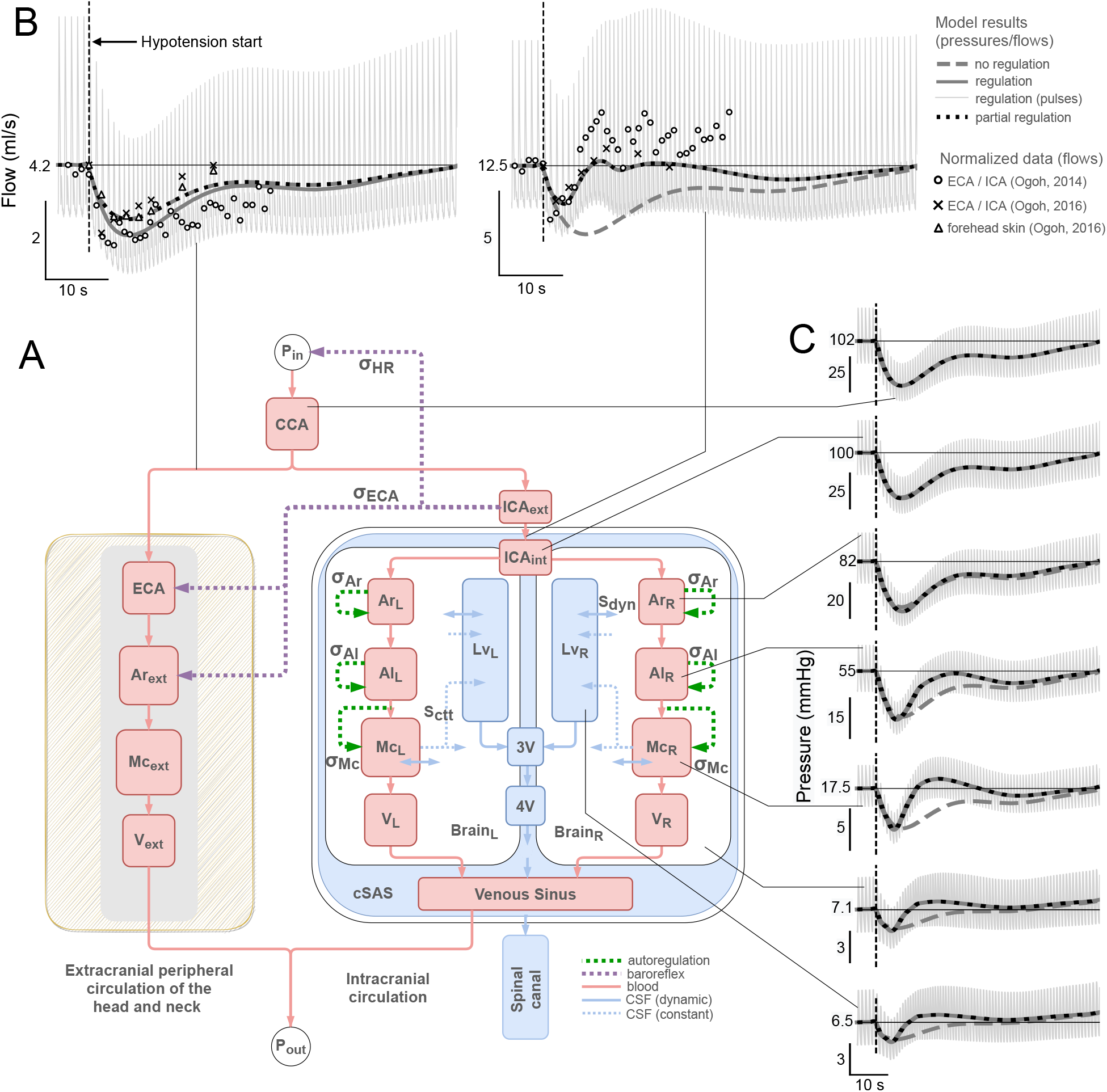
Model overview and results for acute hypotension simulations. **A**: Compartmental model including cerebral, head and neck blood circulations (red), CSF system (blue), biphasic brain parenchyma (white), cerebral autoregulation (green) and baroreflex (purple). The model is based on Linninger et al., 2009^13^ with added compartments and regulation mechanisms. See methods section and Supplementary Section 2 for a detailed explanation. Outline of the intracranial compartments adapted from Linninger model^13^. **B**: Blood flow to the brain (right) and to the peripheral circulation of the head and neck (left) as a function of time, obtained during simulations of acute hypotension in three runs (R_1_ (no regulation): all elements are passive; R_2_ (regulation): cerebral autoregulation, baroreflex control of heart rate and of the peripheral circulation at ECA vascular bed are activated; R_3_ (partial regulation): cerebral autoregulation and heart rate control are activated, but peripheral circulation control is not), as indicated by the top right legend. Horizontal thin lines indicate the model’s baseline levels before the bout of hypotension. The time and amplitudes at different scales are depicted respectively by the horizontal and vertical lines below each graph. Thick curves represent mean flows. Pulsatile flows are only shown for run R_2_. Experimental data^2,3^ superimposed over the model’s response were normalized by the first data points which were aligned to the model’s baselines. **C**: Pressure as a function of time for several compartments, under the same simulation conditions as the flows in **B**.

In order to validate the model, simulations were run and compared to experiments reported in the literature. Especially, published data from experiments of acute hypotension in humans by Ogoh et al.^3,4^ were utilized as the main comparative basis since such experiments recorded the flow at both the ICA and ECA vasculatures. Therefore, we simulated an acute drop in blood pressure in the model by inputting the MAP which was fitted from published data^3^.

The next sections are organized as follows. First, the results of the model in response to an acute drop in blood pressure and the effects of arterial stiffness and of the arterial pressure waveform on the model are presented. In Supplementary Section 3 we included simulations and methods for an acute onset of hypertension, a study of the relations between the intracranial and extracranial blood flows and steady-state analysis of CBF as a function of MAP. These results available in Supplementary Section 3 are also briefly presented and here. We then discuss the results in light of experimental evidence and prior literature. Finally, the model is explained and concluding remarks are given.

## Results

### Relations between CBF and the peripheral circulation of the head and neck

Fig 1 (**B**) shows CBF (right) during acute hypotesion simulations and the blood flow to the ECA vascular bed (left), which models the total peripheral circulation of the head and neck. CBF was reestablished to the baseline level in approximately 7.5 s with a small overshoot (2.24%), while the peripheral blood flow remained suppressed when CA was activated (R_2_ and R_3_). In the model, CBF changed negligibly when regulation at the ECA vascular bed via baroreflex was present (R_2_) in comparison to when it was absent (R_3_). This agrees with recent work suggesting that during acute hypotension the ECA vascular bed does not help reestablishing CBF in healthy young men^4^.

The CBF dynamics in Fig 1 (**B**) is qualitatively in accordance with experiments^3,4,21–23^. The aforementioned experiments reported a return to baseline levels of the blood flow rate and blood flow velocity at the large cerebral arteries from 5 to 17 s after the start of the bout of acute hypotension. The return in 7.5 s of CBF in our model matches well with these times. The overshoot of CBF in the model was 21.6% and 5.9% smaller than those in (Ogoh, 2014)^3^ and (Ogoh, 2016)^4^-control, respectively. The extracranial blood flow dynamics in Fig 1 (**B**) agrees with the dynamics from experiments^3,4^.

Supplementary Table S4 shows the nonlinear statistical relation (NCMI) between CBF and the blood flow to the head and neck. In summary, it measures the similarity of the curves obtained for CBF and for the extracranial blood flow. Their similarity was modest for simulations where CA was turned on (NCMI of 0.37 ± 0.11) and higher (NCMI of 0.55 ± 0.08) when CA was turned off. A more detailed explanation of this statistical analysis is given in Supplementary Section 3.

### Cerebral autoregulation protects the microvasculature

The pressures at several intracranial compartments during acute hypotension simulations are presented in Fig 1 (**C**). When CA was activated (R_2_ and R_3_), the model predicts a faster reestablishment of the pressures to baselines at the brain parenchyma and at the microcirculation than at the arteries and arterioles. These results in Fig 1 (**C**) suggest a mechanistic explanation of the protective role of CA to small vessels and to the brain. Similarly in Supplementary Fig S4, the microcirculation and the brain were protected against a large increase in pressure on the regulated simulations of an acute onset of hypertension.

There is a lack of literature on the effects of acute hypotension to ICP in healthy subjects since such experiments cannot be ethically justified^24^. Therefore, it is difficult to validate our ICP results in Fig 1 (**C**) with clinical data. In severely head-injured patients, the ICP decreased after the cuff deflation by an average of 5 mmHg, returning to baseline levels in 17 s followed by a late overshoot whose peak occurred 55 s after the maneuver^25^. The MAP decreased by on average 19 ± 5 mmHg achieving its minimum 8 ± 7 seconds after the cuff deflation^25^. Here our input MAP decreases by 33.2 mmHg and the nadir occurs 6.9 s after the hypotension start.

Fig 2 (**A**) shows the total cerebrovascular flow resistance and the flow resistances for each of the regulated compartments normalized by the corresponding baseline values. The HR response is presented in **B**. The total cerebrovascular resistance in our model decreased by 27.7%, which is comparable to data from experiments in humans. The total cerebrovascular resistance of 10 healthy subjects in response to a bout of acute hypotension decreased on average by 21.9% on hypocapnia, 24.1% on normocapnia and 21.5% on hypercapnia^21^.

**Figure 2.**
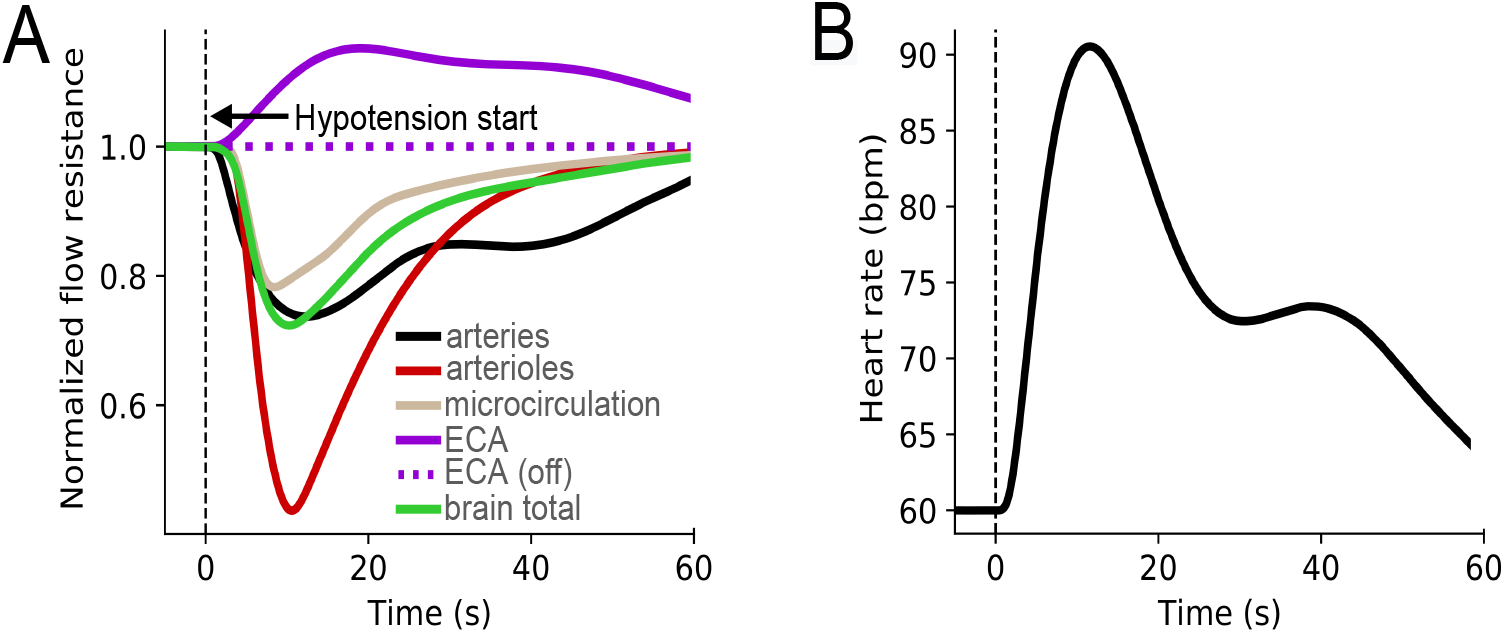
Normalized flow resistances of the regulated compartments and heart rate during acute hypotension simulations. **A**: Resistances of each of the regulated compartments normalized by the baseline values, when the simulation setup included cerebral autoregulation, baroreflex control of heart rate and of the peripheral circulation. The total brain cerebrovascular resistance (green) was calculated by applying principles of resistances in series and in parallel. Since the other regulated simulation run differed only by the ECA regulation, which was turned-off, its results are only shown for the ECA compartment. All other active compartments changed negligibly on those conditions. **B**: Heart rate response during acute hypotension due to baroreflex mechanism. The zeros of the time axes were placed at the hypotension start, but the simulations contained 20 s prior to it to avoid transient effects in the responses.

### CBF in steady-state

Several simulations with different MAPs (MAP kept constant throughout each simulation) were run until steady-state was approximately achieved. See Supplementary Section 3 for a detailed explanation of these simulations. Supplementary Fig S7 shows steady-state CBF as a function of MAP. The overall shape on regulated cases matches with the static autoregulation curve reported in the literature^26,27^. The upper limit of autoregulation of 175 mmHg matches very well with the 175 mmHg from experiments in rats^26^. Our lower limit of 50 mmHg was lower than the experimental 80 mmHg^26^.

### Arterial stiffness and arterial pressure waveform

The effects of the arterial pressure wave and the arterial stiffness in the model are presented in Fig 3. The figure shows the obtained waveforms of the ICP at the brain parenchyma (**B**), CBF, total cerebral venous outflow an the inflow at the spinal cord (**C**), and the calculated change in cerebral blood volume (CBV) at each cardiac cycle (**D**). Accordingly to the input arterial pressures (**A**), the obtained ICPs have increasing peak pressures and amplitudes as age increases. The model predicts ICPs with increasing pulsatilities and CBFs with decreasing pulsatilities as arterial stiffness increases. The change in CBF at each cardiac cycle in Fig 3 (**D**) has a waveform similar to the ICP waveform (**C**) in simulations with normal elastances. This may illustrate that the pulse pressure at the CSF, which is very close to the pulse pressure at the brain parenchyma, is governed by the pulsatile change in CBV and the craniospinal elastance^28^.

**Figure 3.**
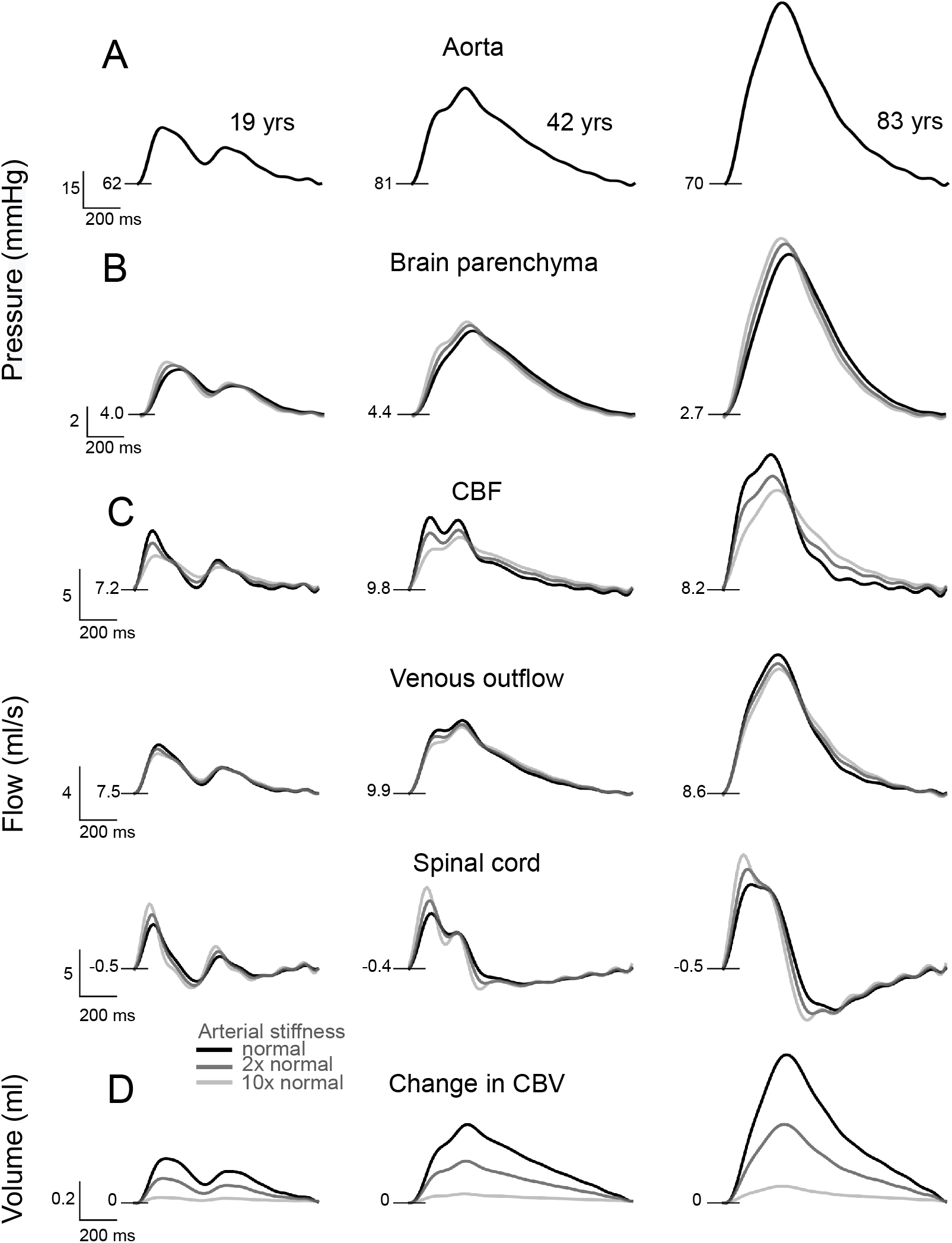
The effects of the arterial pressure waveform and the arterial stiffness on the pressure and flow waveforms in the model. **A**: Aortic pressure waves of individuals aging 19, 42 and 83 years fitted from published data^29^ are set as the pressure at the base of common carotid artery in the model. Three simulations with varying arterial stiffness are shown. **B**: Model’s predictions for the ICP at the brain parenchyma. **C**: Model’s predictions for cerebral blood flow, total cerebral venous outflow and the inflow at the spinal subarachnoid space. **D**: Change in cerebral blood volume at each cardiac cycle.

## Discussion

In this study we intended to build a model including the main aspects of CBF regulation, a good description of the intracranial dynamics and the extracranial vasculature of the head and neck, which potentially acts as a compensatory blood reserve for CBF. The ECA vasculature may be a source of noise for noninvasive technologies that measure the intracranial state^10–12^, therefore a model that includes the peripheral circulation of the head and neck is a first key step to predict how much the superficial tissue influences such devices. The model was validated in comparison to published experimental data from acute hypotension experiments in humans^3,4^.

(Ogoh, 2016)^4^ hypothesised that ECA blood flow may be regulated by a change in perfusion pressure regardless of sympathetic activity. Based on our results we raise the additional hypothesis that the ECA vascular bed may be regulated by sympathetic activity, but the effects of extracranial blood flow regulation on CBF regulation are so small that (Ogoh, 2016)^4^ were not able to measure any significant influence of the ECA vascular bed regulation on CBF regulation.

Understanding how the intracranial and the head and neck blood flows are related is relevant because the extracranial vascular bed is more accessible. For example, if there was a clear dynamic relation between the intracranial and extracranial blood flows, measurements uniquely on the external blood flow could provide an assessment of the intracranial state, since changes in CBF would be perceived at the ECA blood flow. Some studies reported relations between the blood flow at the ICA and ECA vasculatures. For example, after carotid endarterectomy, which reduces the flow resistance to the brain, there was a concomitant reduction in the ECA blood flow and an increase in the ICA blood flow for most of the patients (ECA flow reduced 5.1±48.2%, ICA flow increased 74.9±114.9%)^30^. The opposite behavior may be observed when the ICA flow resistance increases. When the ICA is occluded, the ECA blood flow may increase and the ECA may contribute to CBF trough anastomotic channels^31^.

These studies^30,31^, however, considered longer timescales than the timescale of the acute hypotension experiments. During acute changes in MAP, experimental data and the simulation results suggest that the hemodynamics of the ICA and ECA vascular beds are weakly related in healthy subjects because of different regulatory mechanisms acting on the cerebral vasculature (autoregulation) and possibly at the ECA vasculature (baroreflex). Indeed, tissues presenting intact autoregulation are able to regulate their own blood flow, in a large degree, independently of the flow to tissues in parallel^32^. Conversely, when CA was inactivated in our simulations (analog to impaired CA), the intracranial and extracranial blood flow dynamics became more similar, approximating to a passive dynamics.

We also attempted to predict changes in the intracranial waveforms as a function of arterial stiffness and the arterial waveform because changes in ICP and CBF waveforms have been investigated as indicators for patient condition^28,33,34^. The propagation of the arterial pressure wave along the arterial tree involves complex phenomena such as wave reflections, non-linear elastances of the vessels’ walls and non-Newtonian fluid properties of the blood^35^. An aging vascular system is usually related to an increase in the arterial pulse pressure and in the P2/P1 ratio of the arterial waveform. Studies showed that the flow pulsatility at the middle cerebral artery increases with arterial stiffness^36,37^. The model predicts physiologically plausible increases in the amplitude and in the pressure pulsatility at the brain parenchyma as age and stiffness increase (Fig 3 (**B**)). However, it was expected that the CBF pulsatility also increased, but the model predicted a decrease in CBF pulsatility as arterial stiffness increases (Fig 3 (**C**)). Thus, based on these results and on prior literature^35,38^, we conclude that the model is unsuitable to predict changes in blood flow waveforms as a function of stiffness, as these phenomena depend on distributed parameters (1D and 3D)^35^, while the model is lumped (0D). Despite these limitations, the model predicted an increasing P2/P1 ratio as age increased. Literature suggests that in addition to intracranial compliance, the arterial input contributes to the ICP waveform^39^. Therefore, studies on the ICP wave morphology should also consider the arterial pressure wave as an important factor, as this is a commonly disregarded variable.

## Methods

In this study no experiment in humans nor in animals was performed, therefore no approval from an Institutional Review Board nor from an ethics board for animal research was required.

A model of the intracranial cerebral vasculature divided into compartments representing left and right [arteries, arterioles, microcirculation, veins] and venous sinus which is common to both sides was derived from the Linninger model^13^. The Linninger model^13^ was chosen because it integrated results from extensive magnetic resonance imaging studies^40–43^ with detailed two and three-dimensional mathematical models. It includes the ventricular CSF system divided into left and right lateral ventricles (Lvs), third (3V) and fourth ventricles (4V), cranial subarachnoid space (cSAS) and spinal canal. Lvs are connected to the 3V, which is connected to the 4V. 4V communicates with the cSAS which is connected to the spinal canal. The brain parenchyma is modeled as an incompressible solid cell matrix surrounded by extracellular fluid and is split into right and left hemispheres.

### Extracranial circulation of the head and neck and control mechanisms

The arterial network of the Linninger model^13^ was expanded to include the bifurcation of the common carotid artery (CCA) into the ICA and ECA vasculatures to model the distribution of blood flow to the brain and to the peripheral circulation of the head and neck. The compartment originally denoted by capillaries here is denoted as microcirculation to represent the terminal arterioles, the capillaries and the venules^44^. The venules in the Linninger model were therefore removed. Here, the blood flows from the the CCA tube that bifurcates into the ICA, which supplies the intracranial vessels, and the ECA tube, which supplies the extracranial [arteries, microcirculation and veins]. These compartments model the head and neck peripheral circulation. The total extracranial (ECA + Ar_ext_ + Mc_ext_ + V_ext_) resistance was chosen to enforce a blood flow ratio between cerebral and peripheral circulation in the order of 3, because the head and neck receive about 4% of the cardiac debt, whereas the brain receives approximately 12%^45^. The ICA is divided into the extracranial (ICA_ext_) and intracranial (ICA_int_) segments. Local CA mechanisms acting on the intracranial [arteries, arterioles and microcirculation], and baroreflex control of HR and of peripheral vasculature were added. Fig 1 shows a model overview. The complete model rationale is given in Supplementary Section 2.

### Mathematical formulation

The main mathematical relations for the intracranial dynamics are shown in the Linninger model^13^. Here we summarize the essential equations and focus on added CA and baroreflex. A complete list of the model equations is given in Supplementary Appendix A1.

#### Fluid dynamics

The pressure-induced fluid exchange between two communicating compartments is modelled as a linear equation:

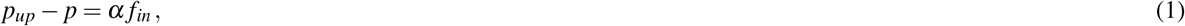

where *p_up_* is the pressure of the upstream compartment, *p* is pressure of the actual compartment and *f_in_* the flow rate entering the compartment. *α* is a constant resistance term for the passive vessels, and is a controlled variable for the regulated vessels. The baseline values for *α*’s were set to reproduce a physiologically valid pressure drop along the vessels and a normal steady-state CBF in the order of 12.5 ml/s.

The compartment’s passive expansion or contraction in response to a change in the transmural pressure is given by:

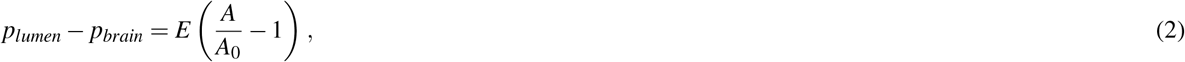

where *E* is the compartment’s wall elastance (stiffness of the vessel), *p_lumen_* is the vessel’s lumen pressure and *p_brain_* is the brain parenchyma pressure on the corresponding side of the vessel.

Finally, the principle of mass conservation is enforced:

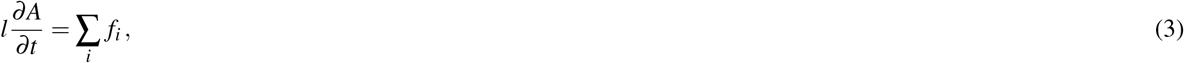

where *f_i_* is the *i*’th flow entering or leaving the compartment. Inflows are taken with positive sign.

#### Cerebral autoregulation

We assume that each compartment’s diameter (*d*) is given by *d* = *d_n_*(1 + *δd*), where *d_n_* is its baseline value and *δd* is the normalized deviation from baseline. Therefore, in the non-regulated case we have *δd* = 0. In the regulated cases, *d* deviates from baseline. If for example the diameter doubles due to autoregulation we have *δd* = 1. For each of the controlled compartments we consider that its diameter deviates from baseline according to a sigmoid function:

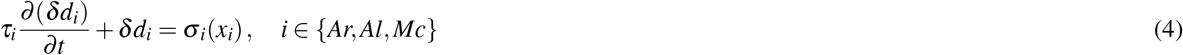

The sigmoid functions based on experimental data^46^ and the parameters are shown in Table 1.

**Table 1.**
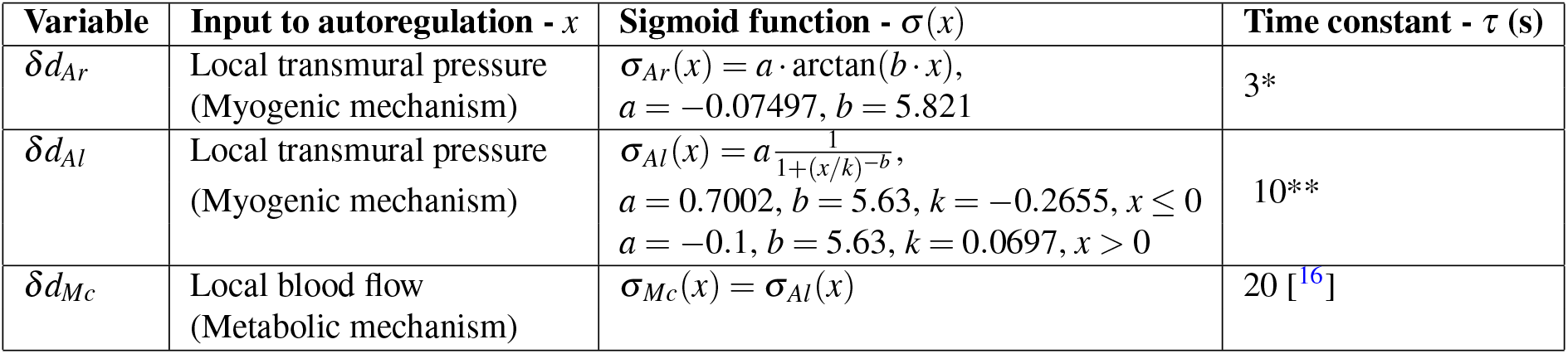
Parameters and their source for autoregulation controlled variables. References are shown within brackets. All the inputs contain an additional pure delay of 1 s for hypotension and 3 s for hypertension simulations due to an online FIR filter designed to estimate their mean values. * is within the range reported in the literature for the myogenic effect^22,47^, ** was chosen as an intermediate value because fast and slow mechanisms act on arterioles. The model’s CBF is highly dependent on the parameters in this Table. In future work they could be tuned to reproduce different autoregulation indexes^23^.

| Variable | Input to autoregulation - $x$ | Sigmoid function - $\sigma(x)$ | Time constant - $\tau$ (s) |
| --- | --- | --- | --- |
| $\delta d_{Ar}$ | Local transmural pressure<br>(Myogenic mechanism) | $\sigma_{Ar}(x) = a \cdot \arctan(b \cdot x)$,<br>$a = -0.07497, b = 5.821$ | 3* |
| $\delta d_{Al}$ | Local transmural pressure<br>(Myogenic mechanism) | $\sigma_{Al}(x) = a \frac{1}{1+(x/k)^{-b}}$,<br>$a = 0.7002, b = 5.63, k = -0.2655, x \leq 0$<br>$a = -0.1, b = 5.63, k = 0.0697, x > 0$ | 10** |
| $\delta d_{Mc}$ | Local blood flow<br>(Metabolic mechanism) | $\sigma_{Mc}(x) = \sigma_{Al}(x)$ | 20 [16] |

This equation is based on the Ursino-Lodi models^15,16^, although here we offer a more complete autoregulation description as we consider that it acts on arteries, arterioles and microcirculation on both sides of the brain, coupled with a detailed model of the intracranial dynamics. The sigmoid functions also differ from Ursino-Lodi models and here we regulate the normalized deviations, not the variables directly. This new idea of controlling only the deviations was necessary to couple autoregulation with the Linninger model while keeping approximately the original baseline levels.

Having *δd* from Eq 4, the vessel’s cross-sectional area at zero transmural pressure follows from:

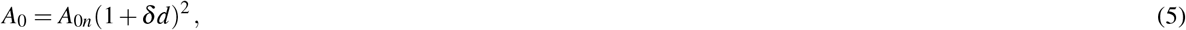

where *A_0n_* is the vessel’s baseline cross-sectional area at zero transmural pressure.

The regulated flow resistance follows from:

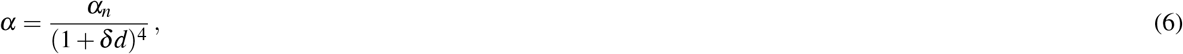

where *α_n_* is the vessel’s baseline flow resistance.

We chose to regulate only the vessel’s resistance and its area at zero transmural pressure, but not the elastance. Changing *A*_0_ and not *E* corresponds to a parallel upward or downward shift of the linear Eq 2. A parallel shift offers a reasonable approximation to the experimental data^48^, although the elastance also had a statistically significant change on constriction and dilation^48^.

#### Baroreflex mechanism

The resistance and cross-sectional area at zero transmural pressure of the ECA and ExAr compartments are controlled by the baroreflex in the same manner as all other controlled vessels (Eqs 4, 5 and 6), trough the shift of their diameter from baseline that follow a sigmoid function (*σ_ECA_*) which was adapted from the Lau-Figueroa model^14^:

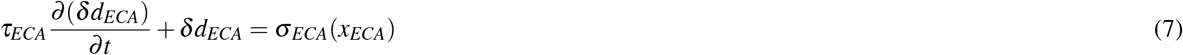

We assume that the normalized shifts from baseline at the ECA and extracranial arteries are equal: *δd_ExAr_* = *δd_ECA_*.

The normalized heart rate 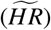 active control by the baroreflex is assumed to be affected both by the sympathetic and parasympathetic tones^14^.

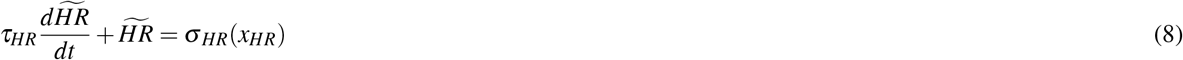

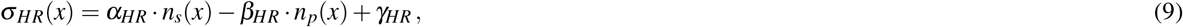

where and *α_HR_* and *β_HR_* are respectively weights for the sympathetic and parasympathetic contributions. *γ_HR_* is a constant that sets the normalized heart rate in baseline conditions equals to 1. The normalized sympathetic and parasympathetic neural activities are given by

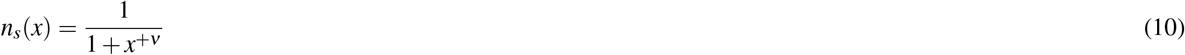

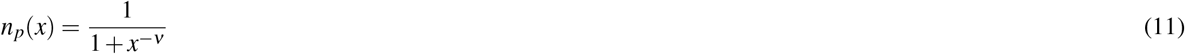

The input *x_HR_* is taken as 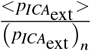, which is the ratio between the mean pressure at the extracranial ICA segment estimated by an online low-pass filter <*p*_*ICA*_ext__> and the extracranial ICA normal pressure (*p*_*ICA*_ext__)_*n*_. Finally, the absolute heart rate *HR* is given by

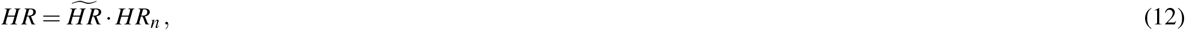

where 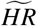 is the normalized heart rate and *HR_n_* is the baseline heart rate. The baroreflex parameters and their source are shown in Table 2.

**Table 2.**
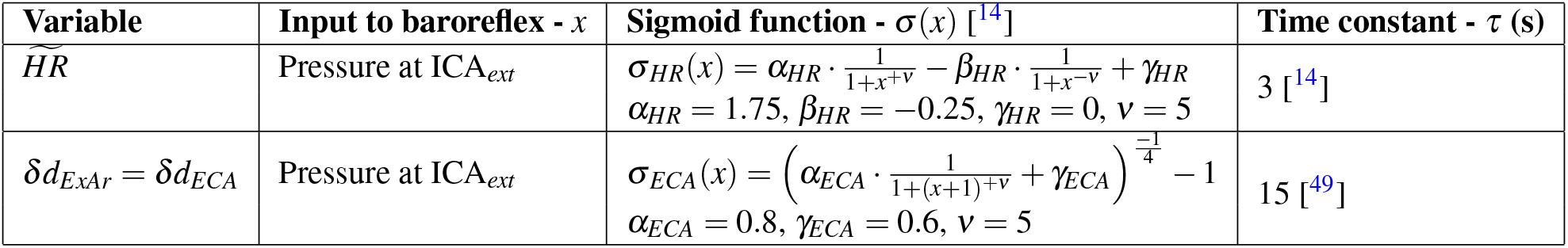
Parameters and their source for baroreflex controlled variables. References are shown within brackets. All the inputs contain an additional pure delay of 1 s for hypotension and 3 s for hypertension simulations due to an online FIR filter designed to estimate their mean values.

The effects of baroreflex on venous capacitance were not included in the model because they affect mostly the splanchnic region^50^, which is unrelated to the intracranial and head and neck blood flow dynamics. The baroreflex effects on heart contractility^50^ were not included because it would require a mechanical model of the heart, which is beyond our goal in this work.

#### Online estimation of mean flows and pressures

The low-pass characteristics of the regulatory mechanisms alone were not able to properly filter the instantaneous values of their inputs. The shape of the mean values suffered alterations and high frequency components passed trough the mechanisms without enough attenuation, producing oscillations in their outputs. A new idea was required so that the model produced physiologically plausible results.

We designed a filter that is integrated on the model to online estimate the mean values of fluxes and pressures that served as inputs to the regulatory mechanisms. We picked a low-pass online finite impulse response (FIR) filter with cutoff frequency of 0.1 Hz and constant group delay (*ϕ*) of 1 s for hypotension and random simulations, which is the same initial delay found in rats kidneys vasodilation^47^, and 3 s for hypertension simulations. The filter order (*N*) follows from 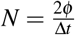, where Δ*t* is the simulation time step in seconds. The filter coefficients were calculated using a Hamming window with length *N* + 1. The filter does not significantly slow the mechanisms because its cutoff frequency is in the order of twice that of the fastest regulatory mechanism (*τ_Ar_* = 3 s). It allows us to know precisely the pure delay (1 s for hypotension and random and 3 s for hypertension simulations) introduced to the regulatory inputs and it preserves the shape of the mean curves. The online filtered variables are denoted as <.> in Supplementary Appendix A1.

In the following we explain the simulations presented here in the main text (acute hypotension; arterial waveforms and stiffness). For the supplementary results (acute onset of hypertension; statistical relation between CBF and the blood flow of the head and neck; CA in steady-state), see explanations on Supplementary Section 3.

### Acute hypotension

To investigate the effects of hypotension in situations corresponding to healthy and impaired conditions, we simulate bouts of acute hypotension by inputting the ABP (*p_init_*) to the model, which drops according to the MAP fitted from published experimental data^3^:

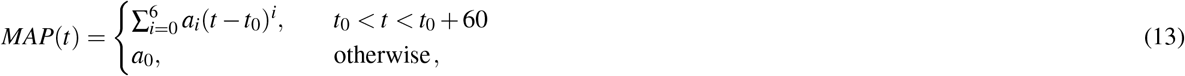

where *t*_0_ = 20 s is the start time of the acute bout of hypotension. *MAP*(*t*) is given in mmHg and *t* in seconds. (*a*_0_,…, *a*_6_) = (102.40, −11.958, 1.4851, −7.4625 · 10^-2^, 1.8273 · 10^-3^, −2.1698 · 10^-5^, 1.0033 · 10^-7^).

The instantaneous ABP at the base of the CCA is given by:

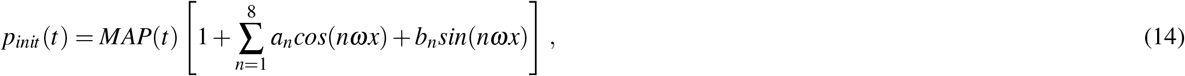

where (*a*_1_,…, *a*_8_) = (−0.0472, −0.0698, −0.0365, −0.0152, −0.0018, +0.0069, +0.0038, +0.0083), (*b*_1_,…, *b*_8_) = (+0.1378, +0.0389, −0.0219, −0.0096, −0.0238, −0.0056, −0.0057, +0.0007). Note that we account for the pulse pressure reduction in the same proportion as the change in MAP.

Using this input we perform three simulations:

(R1) **No regulation**. All regulatory mechanisms are inactivated.
(R2) **Regulation**. CA, baroreflex HR control and baroreflex peripheral circulation control at the ECA and ExAr compartments are activated.
(R3) **Partial regulation**. CA and baroreflex HR control are activated, but peripheral circulation control via baroreflex at the ECA and ExAr is inactivated.

In simulations *R*_2_ and *R*_3_, at the beginning of each heart beat we set *ω* = 2*π · HR*, where *HR* is the last value in Hz of the variable *HR* and *ω* is kept constant until the end of the heart beat to avoid discontinuities on the input pressure. The mean values in Fig 1 were calculated in post-processing using a 5th order low-pass Butterworth filter with cutoff frequency of 0.5 Hz that is applied forwards and backwards.

### Arterial waveforms and arterial stiffness

Three input pressures were fitted from high-fidelity aortic synthesized pressure waves^29^ corresponding to individuals aged 19, 42 and 83 years (Supplementary Table S7). They have increasing arterial stiffness, increasing arterial systolic and pulse pressures^29^. For each subject we considered that P_init_ is the fitted aortic pressure. All regulatory mechanisms were inactivated because we assumed that the individuals are in their baseline conditions. Simulations with varying arterial stiffness were run, considering normal stiffness as those in the original Linninger model. The change in CBV was calculated as 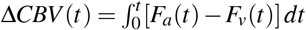, where *F_a_*(*t*) is the total cerebral arterial inflow and *F_v_*(*t*) is the total cerebral venous outflow^28^.

### Limitations

Our model has limitations in addition to the ones already mentioned. The model predicted an overshoot smaller than the experimental data during acute hypotension on regulated simulations. We hypothesize that this is because our model includes the increase in HR, but not the increase in cardiac output during acute hypotension, since cardiac output was experimentally correlated to the increase in ICA blood flow^4^. Modeling the cardiac output was beyond our goal in this work. We did not model the complex regulation of MAP via baroreflex in closed loop. Instead, just the HR and peripheral vasculature are controlled, whereas the ABP is an input to the model, taken from published data. As shown in Supplementary Section 2, we did not calculate the separate contribution of each autoregulation mechanism, but rather we intended to grasp the main factors acting on each compartment depending on the vessels’ sizes. On top of that, we considered that the total flow resistance of the microcirculation, which comprises terminal arterioles, capillaries and venules, is affected by autoregulation. Physiologically, CA only acts up to capillaries. Some limitations are intrinsic to the Linninger model and are described in the seminal work^13^. Long-term mechanisms were not modelled, such as baroreflex resetting in about 48 h^51^.

## Conclusions

We presented a model of cerebral hemodynamics including autoregulation and a good description of the intracranial dynamics and the peripheral circulation of the head and neck, along with baroreflex control of heart rate and of the peripheral vasculature. The first mathematically rigorous explanation to experimental evidence suggesting that the external carotid vascular bed does not help reestablishing cerebral blood flow during acute hypotension in healthy young men is provided. Results suggest that the cerebral and peripheral circulation of head and neck likely become more similar when autoregulation is impaired in comparison to healthy conditions. Simulation results during dynamic changes in mean arterial pressure also provide a possible mechanistic explanation of the protective role of cerebral autoregulation to the microvasculature and to the brain parenchyma. We found that the input arterial pressure wave considerably influences the model’s predictions for the intracranial pressure wave. Having built and validated the model based on experimental data and prior art, we intend in the next steps to predict the noise introduced by the head and neck hemodynamics to noninvasive sensing technologies such as intracranial pressure monitoring by strain gauge and Near-Infrared-based sensors.

## Abbreviations

ABP: Arterial blood pressure
CBF: Cerebral blood flow
CBV: Cerebral blood volume
CA: Cerebral autoregulation
CPP: Cerebral perfusion pressure
MAP: Mean arterial pressure
ICP: Intracranial pressure
CSF: Cerebrospinal fluid
SV: Stroke volume
HR: Heart rate
*P*_xx_: Partial pressure
CCA: Common carotid artery
ICA: Internal carotid artery
ECA: External carotid artery
XX_int_: Intracranial compartment
XX_ext_: Extracranial compartment
Ar: Arteries
Al: Arterioles
Mc: Microcirculation
V: Veins
vSinus: Venous sinus
Lv: Lateral ventricle
3V: Third ventricle
4V: Fourth ventricle
cSAS: Cranial subarachnoid space
br: Brain
*br_solid_*: Brain solid cell matrix
*br_exf_*: Brain extracellular fluid
*p_xx_*: Pressure
<.>: Mean value using a low-pass filter
*xx_n_*: Baseline value of variable *xx*
*xx^L,R^*: Signifying two equations, one for the left and one for the right brain hemisphere
*xx^R^*: Right compartment
*xx^L^*: Left compartment
*f_xx_in__*: Flow into the compartment
*f_xx_out__*: Flow out of the compartment

## Data availability

Model parameters are available in the main text and in Supplementary Appendix A2. On publication raw simulation results and input data will be available at Ambrosio Garcia, Francisco; Lineu Spavieri Junior, Deusdedit; Linninger, Andreas, 2021, “Data for: Cerebral hemodynamics: a mathematical model including autoregulation, baroreflex and extracranial peripheral circulation”, https://doi.org/10.7910/DVN/F5A5NG, Harvard Dataverse, DRAFT VERSION, UNF:6:3Hk7gooXt4vj1TKpDo7VCQ== [fileUNF].

## Acknowledgements

The authors would like to thank Roland Köberle and Thiago Luiz de Russo for helping improve the manuscript and fruitful discussions.

## Author contributions statement

F.A.G. and D.L.S.J. conceived the idea. All authors contributed to the methodology, to the visualization, to the analysis of the results and to the review of the manuscript. F.A.G wrote the main manuscript text. F.A.G. and A.L. developed the software. D.L.S.J. administered the project and acquired funding.

