## Supplementary material for "Cerebral hemodynamics: a mathematical model including autoregulation, baroreflex and extracranial peripheral circulation"

#### 1 Background

##### Cerebral autoregulation

CA is believed to comprise four mechanisms: myogenic, metabolic, endothelial and neurogenic[1,2].

**Myogenic**. The smooth muscle of small cerebral arteries and arterioles contracts when the intralumial pressure increases and relaxes when the pressure decreases[2,3]. This mechanism has the fastest dynamics. In rats cerebral and mesenteric small arteries it started to act with a delay usually less than 250 ms[4]. A median time constant in the order of 6.75 s fitted from eight subjects was found[5]. Note that delay means a dead time during which the mechanisms have no action, while time constant indicates the time after the initial delay needed to achieve 63.2% of the final value in response to a step. This mechanism was included in the model as being pressure-mediated at the arteries and arterioles.

**Metabolic**. Many pathways cause vasomotor changes in small vessels, such as terminal arterioles and capillaries, as a function of tissue metabolism [2,5–8]. For example, high $P_{CO_2}$ and low tissue $P_{O_2}$ cause significant relaxation of cerebral blood vessels[1]. Other contributing factors are tissue concentration of adenosine, lactate, and pH[1]. The time constant is in the order of 20 s[9]. This mechanism was included in the model as being flow-mediated at the microcirculation.

**Endothelial**. Several factors influence the secretion of vasodilators and vasoconstrictors by endothelial cells of brain blood vessels. For example, the secretion of NO contains a shear stress response element[1]. This mechanism is usually modeled as being flow-mediated because shear stress is directly caused by viscous friction of blood and the wall of endothelial cells. It is a slower process with time constant of approximately 60 s[5]. This mechanism was neglected in the model due to its lower contribution when compared to the myogenic one[5].

**Neurogenic**. Neural activity affects the smooth muscle tone of small and medium-sized vessels because cells such as neurons, astrocytes and microglia release neurotransmitters with vasoactive properties[5,7]. For example, neurons secrete NO during activation, which may contribute to increase blood flow[1]. The median time constant was found to be in the order of 8.5 s[5]. This mechanism was not included in the model because it would require an additional input corresponding to the neural activity, which is beyond the goal of our paper

Table S1 summarizes the main aspects of the cerebral autoregulation mechanisms and shows whether or not they are included in our model.

| Mechanism | Main factors | Local of action | Time constant (s) | Model |
|---|---|---|---|---|
| Myogenic | Intralumial pressure[4] | Small arteries and arterioles[3] | 1[10] to 10[5] | Arteries and arterioles |
| Metabolic | $P_{CO_2}$, $P_{O_2}$ [1] | Small vessels [2] | 20[9] | Microcirculation |
| Endothelial | Shear-stress[1,5] | Arteries and arterioles[5] | 60 [5] | Not included |
| Neurogenic | Neural activity | Small and medium-sized vessels[7] | 8.5[5] | Not included |

**Table S1.** Overview of cerebral autoregulation mechanisms and their inclusion or not in the model

##### Baroreflex

The short-term mean arterial pressure (MAP) is controlled mainly by the baroreflex[11]. MAP regulation indirectly influences CBF (indirect regulation). Baroreflex also causes less clear direct vasomotor changes in cerebral blood vessels by modulating sympathetic and parasympathetic activities (direct regulation)[12].

MAP and its rate of change are sensed by baroreceptors, nerve endings that sense stretch, located at the carotid sinuses, especially along the medial wall of the ICA near the common carotid bifurcation[13], and at the aortic arch[14]. The MAP regulates the firing rate of the sensory neurons, which transmit the signals to central integrating centers primarily at the medulla oblongata[15] trough afferent pathways, affecting both sympathetic and parasympathetic tones[15,16]. In addition to the aforementioned baroreceptors, recently it was shown in rats that astrocytes contribute to sympathetic activity, acting as baroreceptors in the brain sensing cerebral perfusion pressure (CPP)[17].

Sympathetic tone modulates the peripheral resistance, venous capacitance mostly in the splanchnic region, heart rate (HR) and heart contractility[16]. The effects of sympathetic activity on different variables are modeled with time constants varying from 3 to 30 s depending on the effector organ[18,19]. The model by Olufsen et al., 2006[20] considered a pure delay of 7 s.

Parasympathetic tone modulates HR, modeled with time constants of 3 to 4 s[18,19], usually without any additional pure delay.

**Indirect regulation.** It corresponds to the indirect contribution of MAP regulation to maintaining CBF[12]. Fig S1 presents the indirect effect of MAP regulation via baroreflex on CBF when MAP drops. We included in the model the control of peripheral vasculature and HR control. The closed-loop mean arterial pressure regulation was not modeled since the pressure at the base of the common carotid artery is the input to our model.

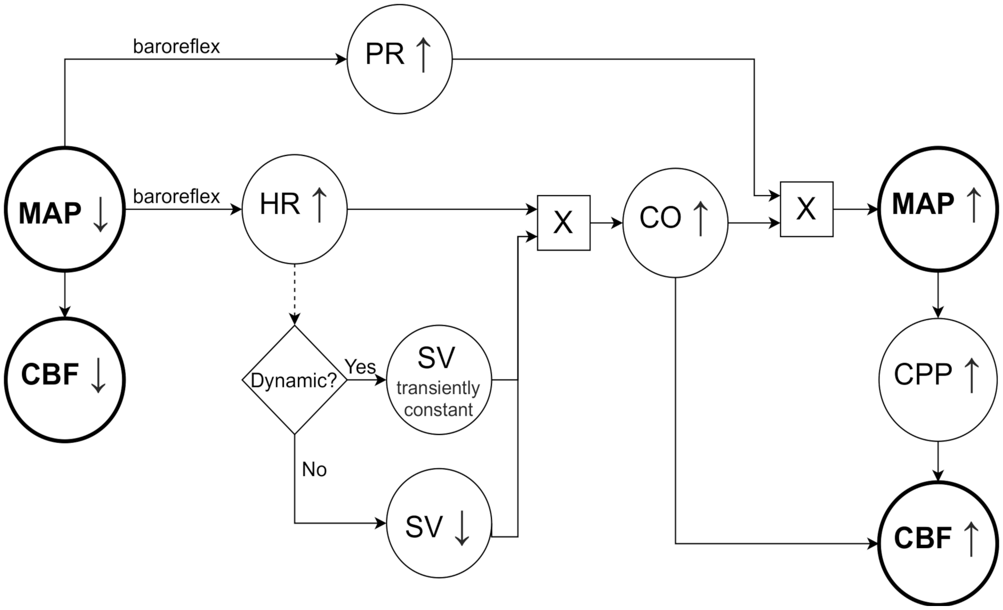


**Figure S1.** MAP regulation via baroreflex and its indirect effect on CBF. When MAP decreases, CBF also decreases. Baroreflex regulation of MAP induces an increase in HR, increase in total peripheral resistance (PR), increase in cardiac contractility and decrease in venous capacitance. These variables affect the stroke volume (SV). In steady-state conditions, an increase in HR is associated to a decrease in SV[21]. Cardiac output (CO) is the product of HR and SV[22]. In steady-state the increase in HR outperforms the decrease in SV, so CO ends up increasing[21]. In specific dynamic situations such as after an acute bout of hypotension caused by a thigh-cuff release, it is thought that SV remains transiently constant, so the increase in CO is proportional to the increase in HR[12]. In both cases MAP increases because it is the product of CO and total peripheral vascular resistance[15], which closes the negative feedback loop. This helps reestablishing CPP, and consequently, CBF, which is also influenced by CO[23].

**Direct regulation.** The direct effect corresponds to alterations in the cerebral vasculature due to the baroreflex and is less clear[24]. In normal conditions, the existing literature reports limited effect in humans, even tough cerebral vessels are innervated with sympathetic nerve fibers[12]. In dynamic situations, (Ogoh, 2016)[23] discusses that sympathetic activity likely plays a role in regulating cerebral vasculature and might regulate ICA vasoconstriction during acute hypotension. In rats it was demonstrated that parasympathetic activity influences cerebrovascular tone[25], especially during acute hypertension[26]. The direct effects of baroreflex on the brain vessels were not included in the model.

#### Literature review on mathematical models

Tables S2 and S3 present our literature review on models of intracranial dynamics, baroreflex and autoregulation. Baroreflex models of a single aspect were excluded.

| Ref | Focus of study | CSF and brain parenchyma systems | PL ICP? | Cerebral autoregulation | BaR | EC |
|---|---|---|---|---|---|---|
| [27,28] | Genesis of the intracranial pressure wave, blood flow in cerebral basal arteries, saline injections and obstruction in the cerebral venous return. | Single capacitance | Yes | Single pressure-mediated | None | Extracranial veins |
| [29] | Arterial hypotension, generation of plateau waves. | Single capacitance | Yes | Pressure-mediated at large vessels, flow-mediated at small vessels | None | Extracranial veins |
| [30] | Generation of plateau waves, acute hypotension, pressure-volume index tests. | Single capacitance | No | Single flow-mediated | None | None |
| [9] | Nonlinear interaction of $CO_2$ reactivity with autoregulation. | Single capacitance | No | Pressure-mediated at large pial arteries, flow-mediated at small pial arteries and their nonlinear interaction with $CO_2$ reactivity | None | Extracranial veins |
| [31] | ICA compression tests, stenosis of ICA and MCA and the role of the Circle of Willis. | Single capacitance | Yes | Same as in [9] | None | ICAs, basilar artery |
| [32] | Relations between cerebrovascular dynamics, ICP, Cushing response and short-term MAP regulation. | Single capacitance | Yes | Flow-mediated and nonlinear interaction with $CO_2$ reactivity | Complete system | Extracranial veins |
| [33] | Simulated MAP and MCA blood flow velocity during postural change from sitting to standing. | Brain resistance | No | Empirical model based on piecewise linear functions | Partial | None |
| [34] | Interactions between cerebral autoregulation and brain gas exchange during carotid artery compression, short/long-term arterial hypotension. | Single intracranial compliance, interstitial and intracellular spaces have constant volumes | Yes | Flow-mediated at arteries and effect of $CO_2$ | Complete system | Neck arteries and veins |
| [35] | Predicted intracranial pressure gradients, blood and CSF flows in normal and in communicating hydrocephalus conditions. | Biphasic brain parenchyma, lateral, third and fourth ventricles, cSAS, spinal cord | Yes | None | None | Jugular veins are boundary condition |
| [36] | Studied the effect of regulatory asymmetry on blood flow and predicted the interhemispheric steal effect. | Single capacitance, left and right cSASs and cerebral aqueduct are resistances | Yes | Pressure-mediated and $CO_2$-mediated acting on single element | None | ICAs and venous outflow compliances |
| [37] | Coupling previous model of cerebrovascular bed[38] with autoregulation models. | None | No | Tested empirical model based on polynomial functions, a simple model and an optimal control model | None | None |
| [5] | Cerebral autoregulation and neurovascular coupling in response to squat-stand maneuvers and visual stimulation. | None | No | Myogenic, metabolic, endothelial, neurogenic | None | None |
| [39] | Contribution of myogenic, shear-stress based and metabolic mechanisms with a theoretical model. | None | No | Myogenic, metabolic, shear-stress based | None | None |

**Table S2.** Overview of intracranial dynamics, baroreflex and cerebral autoregulation models - part 1. Ref: Reference. PL: Pulsatile. ICP: intracranial pressure. BaR: baroreflex. EC: Extracranial circulation. ICA: Internal carotid artery. MCA: Middle cerebral artery. MAP: Mean arterial pressure. CSF: Cerebrospinal fluid. cSAS: Cranial subarachnoid space.

| Ref | Focus of study | CSF and brain parenchyma systems | PL ICP? | Cerebral autoregulation | BaR | EC |
|---|---|---|---|---|---|---|
| 40 | Predicted the role of the head and neck peripheral veins in regional blood redistribution on Earth and on microgravity during posture change. | Single capacitance | No | Single flow-mediated | None | Detailed description of head and neck arteries and veins |
| 19 | Simulated the short-term pressure regulation during the tilt test. | None | No | None | Complete system | 3D model of left and right [subclavians, CCAs bifurcating into ICAs and ECAs] |
| 18 | Effect of baroreflex during hemorrhage in the abdominal aorta artery and on cerebral aneurism in the presence of a regurgitating aortic valve. | None | No | None | Complete system | All main arteries |
| 41 | Studied the impact of bilateral traverse sinus stenosis on cerebral venous flow and CSF dynamics, as well as treatment strategies. | Same as in the Linninger model[35] | Yes | None | None | Detailed description of head and neck veins |
| 42 | Predicted subject-specific flows and pressures in the ventricular system and subarachnoid space. | Entire cSAS and ventricular system in 3D | Yes | None | None | None |
| 43 | Simulated orthostatic stress tests. | None | No | None | Complete system, including cardiopulmonary reflex | None |
| 44 | Studied the cardiovascular response to centrifugation and lower body ergometer exercise. | None | No | None | Complete system, including cardiopulmonary reflex | None |

**Table S3.** Overview of intracranial dynamics, baroreflex and cerebral autoregulation models - part 2. Ref: Reference. PL: Pulsatile. ICP: intracranial pressure. BaR: baroreflex. EC: Extracranial circulation. CCA: Common carotid artery. ICA: Internal carotid artery. ECA: External carotid artery. CSF: Cerebrospinal fluid. cSAS: Cranial subarachnoid space.

References[34, 36, 40] have shown to be the models closest to our goal here in this paper. Lau-Figueroa[34] included the MAP regulation via baroreflex in closed loop and CA only at the arteries. Although the model contains capacitances of the arteries and veins of the neck, their contribution to CBF was not addressed. The compliance of the intracranial space was lumped to a single capacitance.

Piechnik et al., 2011[36] included more than one element on the cerebrospinal fluid (CSF) system but the overall compliance of CSF space was lumped to a single capacitance. They included the myogenic and $CO_2$ reactivity mechanisms on a single side of the brain to study the impact of the regulatory asymmetry.

More recent work[40] offers a detailed description of the head and neck circulations. Their cardiovascular model is coupled with a simple description of the intracranial dynamics[30].

### 2 Model rationale

#### The Linninger model of intracranial dynamics

In this section the Linninger model[35] is briefly explained.

##### *Cerebral blood flow*

In the Linninger model[35] the brain is supplied by a single carotid compartment which bifurcates into right and left cerebral arteries, arterioles, capillaries, venules and veins which converge to the venous sinus. Finally the blood flows to the jugular veins which are a boundary condition consisting of a constant venous pressure ($p_{out}$).

##### *CSF system*

The CSF production occurs at the choroid plexuses which consist of capillaries and connective tissue separated from the ventricles by epithelial cells[45]. CSF production is modelled as a constant term of fluid exchange between the microcirculation and lateral ventricles. The model also accounts for fluid exchange between the brain extracellular fluid and the lateral ventricles.

After passing trough the third and fourth ventricles and the cranial subarachnoid space (cSAS), the CSF is reabsorbed by the venous sinus according to a reabsorption resistance. The cSAS is communicating to the spinal canal, which is free to expand, contrarily to the intracranial compartments whose total volume is maintained constant by enforcing the Monro-Kellie doctrine separately at each side of the skull.

##### *Brain parenchyma*

The brain parenchyma is modeled as a solid cell matrix with constant volume surrounded by a extracellular fluid compartment which may expand, and is split into right and left hemispheres. The extracellular fluid may seep from the capillary bed into the brain in both directions according to the pressure gradient and an exchange resistance. Here it is important to remark that the incompressible cell matrix is a strong assumption that cannot capture for example plastic deformations on the brain tissue[35].

#### Arterial expansion and inclusion of control mechanisms

We added or modified the following elements to the Linninger model:

##### *Extracranial ($ICA_{ext}$) and intracranial ($ICA_{int}$) internal carotid compartments*

Approximately 3/4 of the brain blood supply comes from the internal carotid arteries and 1/4 from the vertebral arteries[24]. Here the flow at the intracranial ICA compartment is intended to mimic the total CBF.

The total ICA length is about 17.7 cm[46] whereas its extracranial part is about 8.6 cm[47]. We model the brain supply as two compartments in series: extracranial ICA ($ICA_{ext}$) and intracranial ICA ($ICA_{int}$) segments with total volume of 5.19 cm³ ($V_0$), which is equal to the total volume of the ICAs and vertebral arteries[48]. We arbitrarily enforced the $ICA_{ext}$ area at rest to be the double of the $ICA_{int}$ area and the elastance to be the half.

Physiologically, the vertebral arteries, whose volume was included in the ICA tubes, branch from the subclavian arteries, not from the CCA. The $ICA_{int}$ is included in the Monro-Kellie equations. These compartments substituted the original single carotid compartment.

##### *External carotid (ECA), extracranial [arteries ($Ar_{ext}$), microcirculation ($Mc_{ext}$), veins ($V_{ext}$)] compartments*

At baseline levels the average ICA blood flow is in the order of 1.5 times the ECA blood flow[49]. The head and neck receive about 4% of the cardiac debt, whereas the brain receives the triple, in the order of 12% of the cardiac debt[48]. Therefore we chose the total extracranial (ECA + $Ar_{ext}$ + $Mc_{ext}$ + $V_{ext}$) resistance to enforce a blood flow ratio between cerebral and peripheral circulation in the order of 3, instead of a blood flow ratio of 1.5.

Assuming a total blood volume of 4.9 l[50], the total volume at rest of the extracranial compartments was calculated to be 239.86 ml, because the head and neck contain about 4.895% of the total blood volume[48]. The ECA length of 6.10 cm was taken from[48], and the ECA area was $2\pi r_{ECA}^2$, where $r_{ECA} = 0.2265$ cm is the ECA internal radius[48]. The total length of the extracranial compartments of 20 cm was chosen arbitrarily.

###### ***Common carotid compartment (CCA)***
The input to our model is the arterial blood pressure (ABP) at the base of the common carotid, which splits into the ICA and ECA arteries. Its length was taken as the average of the right and left common carotids lengths[48] and its area was $2\pi r_{CCA}^2$ where $r_{CCA} = 0.3029$ cm is the common carotid lumen radius[48].

###### ***Microcirculation (MC)***
The Linninger model contains a compartment called Venules (Vl) and one called Capillaries (Cp). The volume of Cp in the Linninger model[35] was taken from the microcirculation tube in Zagzoule-Vergnes model[51], which already includes the volume of the terminal arterioles, the capillaries and the venules. Therefore we decided to remove the Vl compartment and rename the Cp compartment to Microcirculation.

###### ***Autoregulation mechanisms***
CA takes place most notably from the small cerebral arteries[2] up to small arterioles and capillaries[6]. The percentual changes in vessels diameters and their time constants are highly heterogeneous depending on the brain region and vessel sizes[52]. Instead of calculating the separate contribution of each autoregulation mechanism to each compartment, we developed a simpler model which is intended to grasp the main characteristics of CA in function of the vessels' sizes[53]. We therefore chose different input variables and parameters to each controlled compartment. Our autoregulation does not consider the neurogenic mechanism because we did not add an input to the model corresponding to the neural activity.

Usually the large cerebral arteries (ICA, middle and vertebral) are modeled as passive elements[5,9]. Similarly, we assume that the ICA is not regulated. This assumption is supported by experiments[54], which reported that the mean diameter of large cerebral arteries changed less than 4% in patients during craniotomy under altered MAP and end tidal carbon dioxide, whereas smaller arterial diameters changed more than 20%. Despite this, the ICA vasculature may be regulated by sympathetic tone[23], but the evidence is limited.

For the intracranial arteries, arterioles and microcirculation, we consider that each compartment is self-regulated, which is in accordance with the local action of the myogenic and shear stress-based mechanisms. Physiologically this local action does not hold completely true. For example, the metabolic mechanism may act upstream with the diffusion of $CO_2$ from the microenvironment to arterioles[5].

The myogenic mechanism acts on small arteries and arterioles[3], and when compared only to the endothelial shear stress-based mechanism, it highly dominates both in short-time regulation and in steady state[5]. We assume in this manner that the input to the autoregulation of both the arterial and arteriolar compartments is the local transmural pressure. Physiologically, both the pressure and flow contribute to the regulation of arterioles, with the pressure shown to be more important in cats mesentery[55]. When we used the transmural pressure as the input to the arterioles autoregulation, our simulations showed results more close to what we expected to be physiologically plausible.

Instead of calculating the smooth muscle tension[5], we consider that the input variable passes directly trough a sigmoidal function followed by a low-pass filter, similar to the simple Ursino et al. model[30]. To account for heterogeneous diameter changes along different vessel sizes, we considered different sigmoid functions based on data from experiments in cats[56]: $\sigma_{Ar}(x)$ is used for the arteries compartment and $\sigma_{Al}(x)$ which is used both for the arterioles and the microcirculation compartments. We make the distinction that the input for the arteriolar compartment is the transmural pressure while for the microcirculation it is the blood flow, to account for the metabolic mechanism on smaller vessels. They are shown in Fig S2 (**A**) and (**B**). For the arterioles sigmoid we had to find a compromise between the agreement to data[56] and physiological agreement of the results with experiments in humans (e.g.[5,23,49,57]), so that the negative part of our $\sigma_{Al}(x)$ is slightly steeper than a more detailed model by Ursino-Lodi[9]. Its lower limit in the positive part was taken from the literature[39].

During slight hypotension or hypertension, the diameter of the arteries change the most, while the arterioles remain almost unchanged. When MAP continues decreasing or increasing, the arteries' response saturate and the response of the arterioles increases dramatically[9], which is a behavior comparable to a sigmoid containing an initial threshold. The response of cerebral vessels to hypertension is lower than to hypotension[30].

We used the same sigmoid for both the arterioles and microcirculation, in which the terminal arterioles are comprised, because large arterioles and small arterioles display similar activation functions on a theoretical model[39]. We do not account for the late dilation of the arterioles during extreme hypertension[56]. The upper and lower limits of the arteries sigmoid in Fig S2 (**A**) match very well with the diameter dilation and constriction in the order of 11% of the brachial artery in response to nitroglycerin and norepinephrine, respectively[58].

###### ***Baroreflex mechanism***
Cerebral blood vessels are densely innervated with both sympathetic and parasympathetic nerve fibers[1]. The direct effect of sympathetic tone on cerebral vasculature is still not completely understood and varies highly over mammals[59]. In humans it is thought that baroreflex plays an important role in protecting the blood-brain barrier during acute hypertension, but in normal

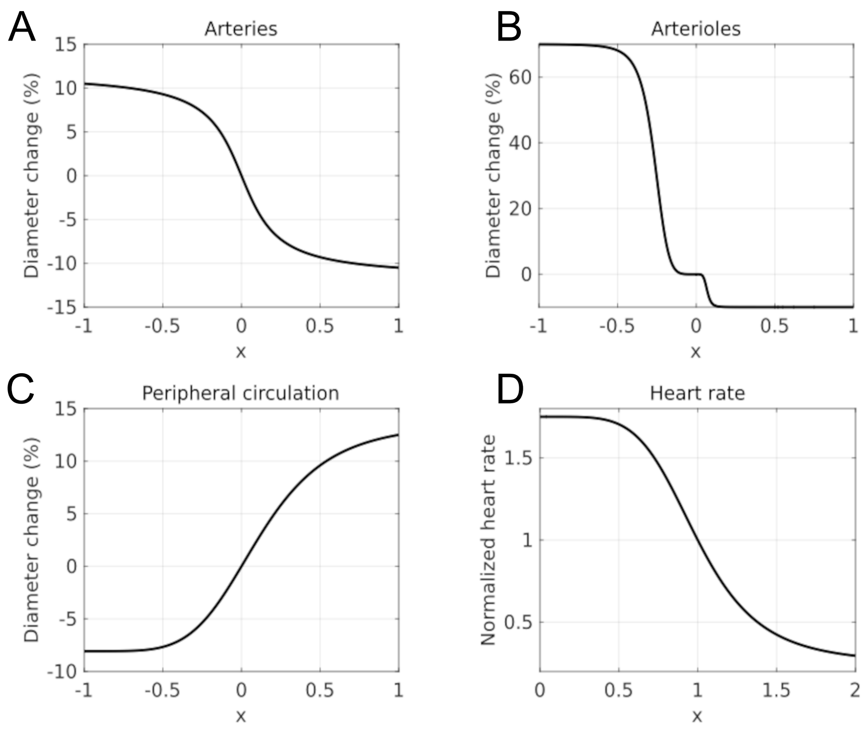


**Figure S2.** Sigmoid functions for each of the regulated elements. **A**: $\sigma_{Ar}(x)$, used for the arteries. **B**: $\sigma_{Al}(x) = \sigma_{Mc}(x)$ , used for the arterioles and microcirculation. **C**: $\sigma_{ECA}(x)$, used for the external carotid and extracranial arteries, adapted from Lau-Figueroa model[19]. **D**: $\sigma_{HR}(x)$, used for the heart rate, taken from Lau-Figueroa model[19]. The central $x$ corresponds to the baseline level for each of the mechanisms inputs. Note that the central $x$ for $\sigma_{HR}(x)$ is 1, whereas for the others is 0 because of different definitions of the inputs. See Eqs 39, 46, 53, 3 and 14 of Appendix A1.

conditions it has little effect[1,12]. We decided to neglect the direct effect of both sympathetic and parasympathetic tones on the cerebral vessels.

We consider that the baroreflex acts only on the diameter of the [ECA and ExAr] and at the HR. We compare the simulations in which the peripheral circulation of the head and neck is and is not controlled by sympathetic tone, because there is limited evidence supporting both cases, although recent work suggests that the ECA vascular bed is passive[23].

The ABP at the base of the common carotid compartment is an input to our model, which in terms of MAP is in open loop. In this way we did not model the complex MAP regulation in closed loop as other excellent models (e.g.[18]). In future work we may close the MAP loop including a mechanical model of the heart. The sigmoid functions used for the peripheral diameter change and HR, which were adapted from the Lau-Figueroa[19] model, are shown in Fig S2 **(C)** and **(D)**.

#### 3 Supplementary results

##### Acute onset of hypertension

We also performed the three simulations described on the main text ($R_1$: **No regulation**, $R_2$: **Regulation** and $R_3$: **Partial regulation**) during an acute onset of hypertension. The input MAP was taken from published data of experiments which induced hypertension in rats[60]. In order to make the input pressure more physiologically plausible to what is expected in humans, we re-scaled the data to set the baseline level equals to 102.40 mmHg and a maximum MAP of approximately 150 mmHg in the end of the hypertensive onset.

Fig S3 shows the obtained CBF and the blood flow to the extracranial vascular bed of the head and neck. Fig S4 presents the pressures at several compartments. The normalized flow resistances for each regulated compartment are in Fig S5.

The mean values in Figs S3 and S4 were calculated in post-processing using a 5th order low-pass Butterworth filter with cutoff frequency of 0.1 Hz that is applied forwards and backwards.

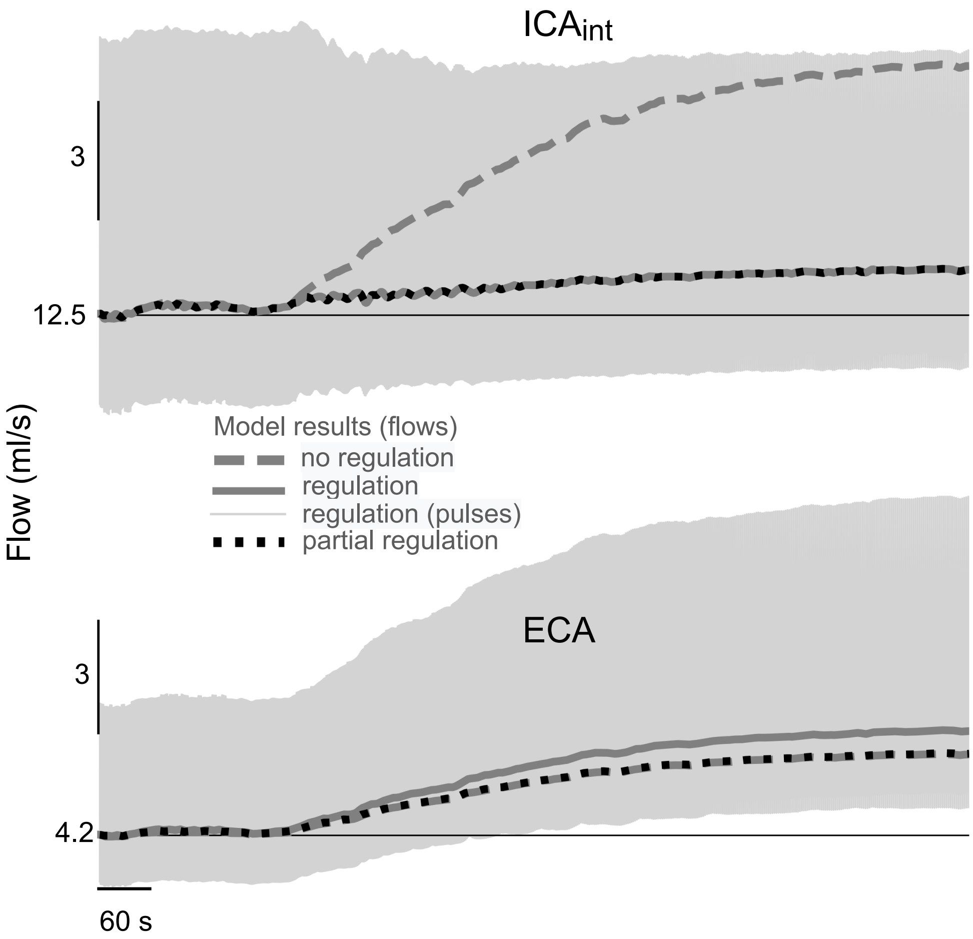


**Figure S3.** Blood flow to the brain (top) and to the peripheral circulation of the head and neck (bottom) as a function of time, obtained during simulations of acute onset of hypertension in three runs ($R_1$ (no regulation): all elements are passive; $R_2$ (regulation): cerebral autoregulation, baroreflex control of heart rate and of the peripheral circulation at ECA vascular bed are activated; $R_3$ (partial regulation): cerebral autoregulation and heart rate control are activated, but peripheral circulation control is not), as indicated by the legend. Horizontal thin lines indicate the model's baseline levels before the bout of hypertension. The time and amplitudes at different scales are depicted by the horizontal and vertical lines below/above each graph, respectively. Thick curves represent mean flows. Pulsatile flows are only shown for run $R_2$.

##### Statistical relation between CBF and the extracranial blood flow of the head and neck

A main goal of this model is to predict in future work how much noise the superficial tissue introduces to noninvasive sensing technologies. As a first step in this direction, we estimated trough the model how the CBF and the superficial blood flow of the head and neck are related in a statistical sense.

We performed the following three extra simulations in addition to the hypotension and hypertension ones for each of the two cases: **Regulated**: CA and HR control are activated; **Non-regulated**: all control mechanisms are inactivated. The ECA vascular bed was considered passive in both cases in order to match with the discussions in (Ogoh,2016)[23].

- 3 simulations inputting random MAPs that follow random walks starting at 102.40 mmHg and whose increments are sampled from a Gaussian distribution with zero mean and standard deviation of 0.15 mmHg.

We then estimated the normalized conditional mutual information (NCMI $\in [0,1]$)[61] of the mean intracranial ($< f_{ICA} >=$ CBF) and extracranial ($< f_{ECA} >$) blood flows, conditioned on the mean input pressure ($< p_{init} >$). The mutual information is a statistical measure of non-linear dependency between variables[62,63]. The conditioning on $< p_{init} >$ was chosen as an attempt to eliminate the effect of the arterial pressure on both the intracranial and extracranial blood flows, as depicted on Fig S6.

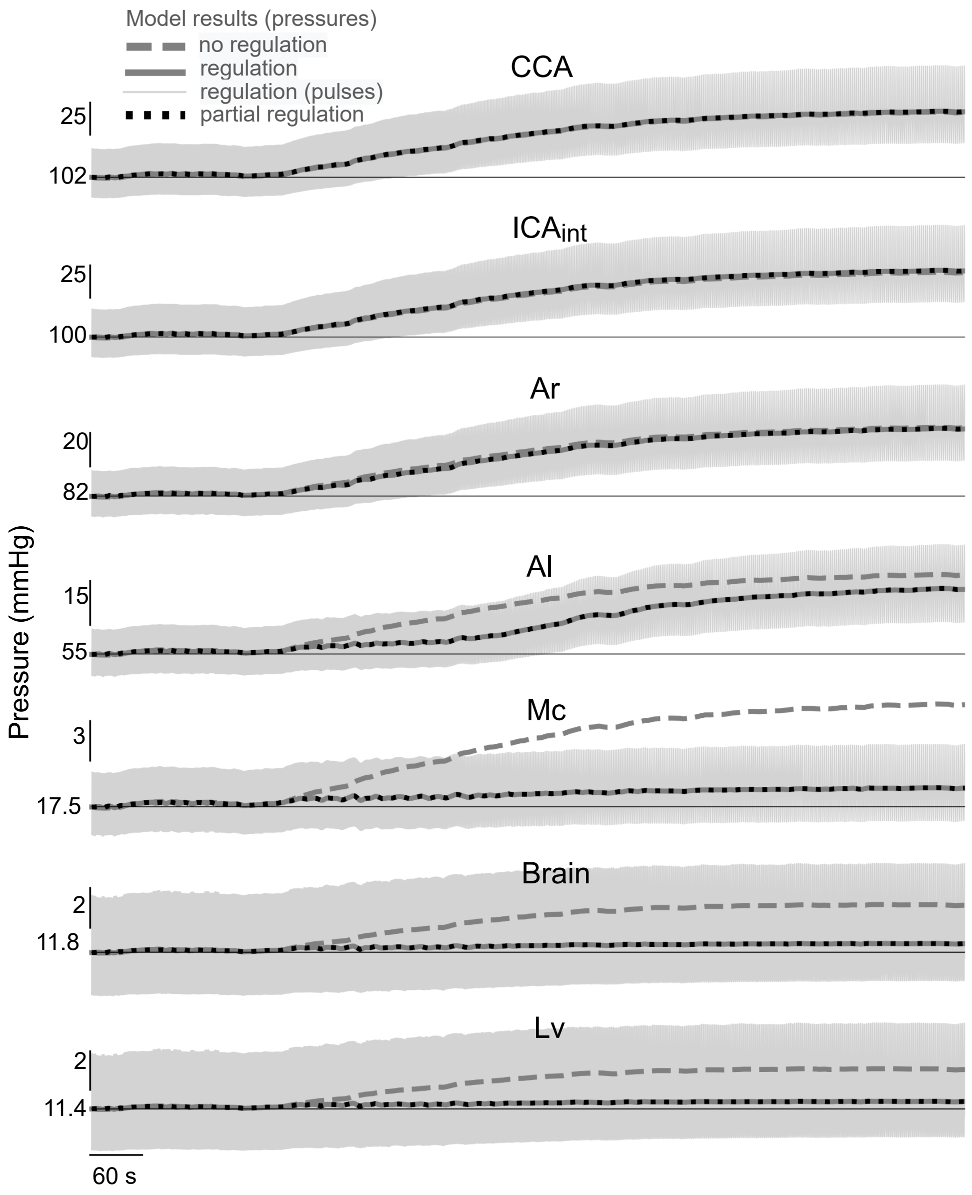


**Figure S4.** Pressures at several compartments obtained at same simulation conditions of acute onset of hypertension as in Fig S3.

$$\text{NCMI}(< f_{ICA} >, < f_{ECA} > | < p_{init} >) = \frac{I(< f_{ICA} >, < f_{ECA} > | < p_{init} >)}{\min\{H(< f_{ICA} > | < p_{init} >), H(< f_{ECA} > | < p_{init} >)\}}, \quad (1)$$

where $I(A,B|C)$ is the conditional mutual information between $A$ and $B$, conditioned on $C$, and $H(A|C)$ is the entropy of $A$ conditioned on $C$. The NCMI was calculated using entropies estimated by the finite sample correction method[64] in Python v.3.6 based on the MDEntropy library[65]. The estimated NCMI at each simulation is shown in Table S4. The mean flows and pressures were estimated using 5th order butterworth low-pass filters that are applied in both directions with cutoff frequencies of 0.5 Hz for the hypotension and random simulations, and 0.1 Hz for hypertension.

The obtained NCMI between CBF and the extracranial blood flow conditioned on the arterial pressure was modest on the regulated case (0.37 ± 0.11). It was particularly low for the hypertension simulation (0.23) and moderate for the hypotension

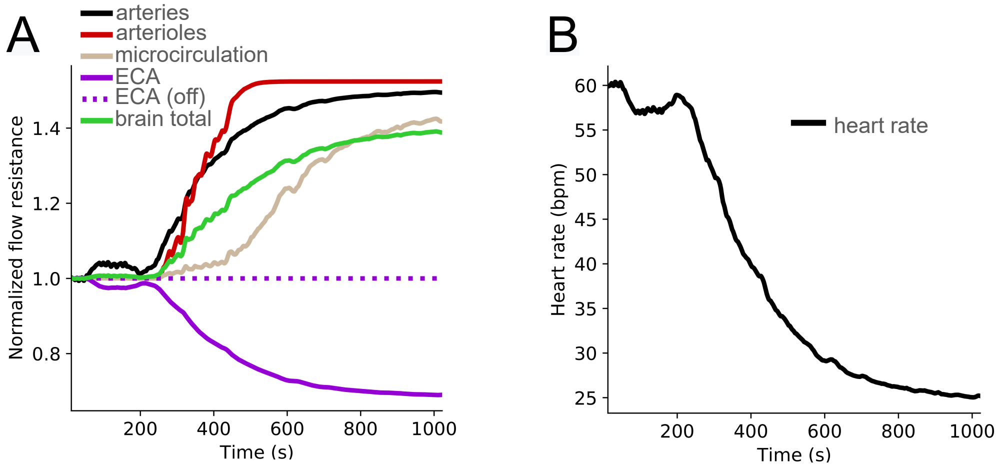


**Figure S5.** Normalized flow resistances of the regulated compartments and heart rate during acute onset of hypertension simulations. **A**: Resistances of each of the regulated compartments normalized by the baseline values, when the simulation setup included cerebral autoregulation, baroreflex control of heart rate and of the peripheral circulation. The total brain cerebrovascular resistance (green) was calculated by applying principles of resistances in series and in parallel. Since the other regulated simulation run differed only by the ECA regulation, which was turned-off, its results are only shown for the ECA compartment. All other active compartments changed negligibly on those conditions. **B**: Heart rate response during acute onset of hypertension due to baroreflex mechanism.

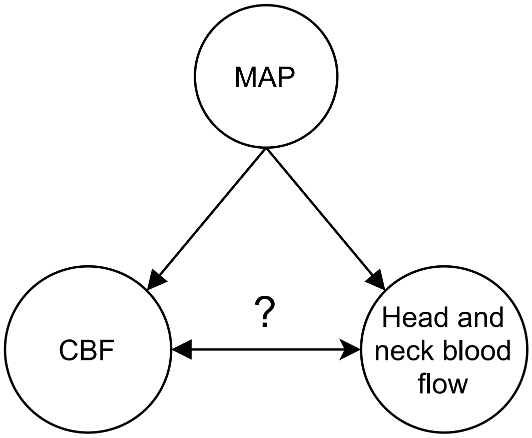


**Figure S6.** Simultaneous effect of mean arterial pressure on intracranial and extracranial circulations. Since our interest lies on the relations between CBF and the peripheral blood flow of the head and neck, highlighted with a question mark, the statistical analysis (NCMI) was conditioned on MAP.

| | **NCMI**$(< f_{ICA} >, < f_{ECA} > \mid < p_{init} >)$ | |
|---|---|---|
| **Simulation** | **Regulated** | **Non-regulated** |
| Hypotension | 0.56 | 0.60 |
| Hypertension | 0.23 | 0.68 |
| Random walk 1 | 0.33 | 0.50 |
| Random walk 2 | 0.35 | 0.50 |
| Random walk 3 | 0.36 | 0.49 |
| Average± SD | $0.37 \pm 0.11$ | $0.55 \pm 0.08$ |

**Table S4.** Normalized conditional mutual information between CBF and the extracranial blood flow of the head and neck, conditioned on the input pressure.

(0.56). Note that the NCMI is a symmetric measure. The average NCMI was moderate (0.55 ± 0.08) on the non-regulated case.

##### Static autoregulation analysis

In order to analyse the proposed autoregulation mechanisms in steady-state, we performed simulations with various MAPs. MAP was maintained constant with zero pulse pressure throughout each simulation. We considered three situations:

(R1) **No regulation**. All regulatory mechanisms are inactivated.

(R2) **Regulation**. CA and peripheral circulation control at the ECA and $Ar_{ext}$ via baroreflex are activated, but HR control is inactivated.

(R3) **Partial regulation**. CA is activated, but HR and peripheral circulation control via baroreflex are inactivated.

To achieve steady-state the system is run for about 5 times the largest time constant ($\tau_{Mc} = 20$ s). The pressure at the brain parenchyma was kept constant in the order of 7.1 mmHg to match the experimental setup [56] and the analysis done in Ursino-Lodi model[9]. To do so we added a term in Eq 77 connecting both brain hemispheres to a compartment with infinite capacitance and constant pressure trough a low flow resistance.

Fig S7 shows the steady-state CBF in function of MAP for the three situations. We plot the range of normal CBF as baseline CBF ±20%[66]. The obtained upper limit of autoregulation was 175 mmHg while the lower limit was 50 mmHg.

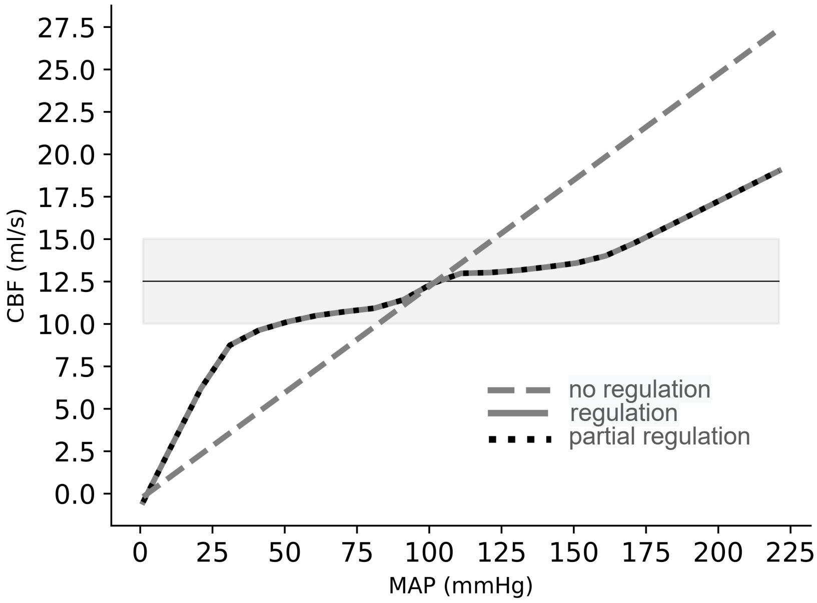


**Figure S7.** Static autoregulation curve - cerebral blood flow as a function of mean arterial pressure in steady-state. Autoregulation curves are drawn for three simulations. $R_1$ (no regulation): All regulatory mechanisms are inactivated; $R_2$ (regulation): CA and peripheral circulation control at ECA vascular bed are activated; $R_3$ (partial regulation): Only CA is activated. Normal CBF range is shown as baseline CBF ±20%, which is an arbitrary definition[66].

### Appendix

#### A1. Equations

**Heart rate**

$$\tau_{HR}\frac{d\widetilde{HR}}{dt} + \widetilde{HR} = \sigma_{HR}(x_{HR}), \quad \text{normalized HR control via baroreflex} \tag{2}$$

$$x_{HR} = \frac{< p_{ICA_{\text{ext}}} >}{(p_{ICA_{\text{ext}}})_n}, \quad \text{input to baroreflex: pressure at } \text{ICA}_{\text{ext}} \tag{3}$$

$$HR = \widetilde{HR} \cdot HR_n \,, \quad \text{absolute value of heart rate} \tag{4}$$

**Input pressure**

$$p_{init} = MAP(t) \left[ 1 + \sum_n a_n cos(n\omega x) + b_n sin(n\omega x) \right] \,, \quad \text{pressure at base of CCA} \tag{5}$$

$$\omega = 2\pi HR \,, \quad \text{beat-to-beat fundamental frequency} \tag{6}$$

**Common carotid artery (CCA)**

$$l_{CCA} \frac{\partial A_{CCA}}{\partial t} = f_{CCA\,in} - f_{CCA\,out} \,, \quad \text{continuity} \tag{7}$$

$$p_{init} - p_{CCA} = \alpha_{CCA} f_{CCA\,in} \,, \quad \text{momentum} \tag{8}$$

$$p_{CCA} - 0 = E_{CCA} \left( \frac{A_{CCA}}{A_{CCA0}} - 1 \right) \,, \quad \text{distensibility} \tag{9}$$

**External carotid artery (ECA)**

$$l_{ECA} \frac{\partial A_{ECA}}{\partial t} = f_{ECA\,in} - f_{ECA\,out} \,, \quad \text{continuity} \tag{10}$$

$$p_{CCA} - p_{ECA} = \alpha_{ECA} f_{ECA\,in} \,, \quad \text{momentum} \tag{11}$$

$$p_{ECA} - 0 = E_{ECA} \left( \frac{A_{ECA}}{A_{ECA0}} - 1 \right) \,, \quad \text{distensibility} \tag{12}$$

$$\begin{cases} \tau_{ECA} \frac{\partial(\delta d_{ECA})}{\partial t} + \delta d_{ECA} = \sigma_{ECA}(x_{ECA}) \,, \text{ case in which baroreflex acts on ECA} \\ \delta d_{ECA} = 0 \,, \text{case in which baroreflex does not act on ECA} \end{cases} \tag{13}$$

$$x_{ECA} = \frac{< p_{ICA_{\text{ext}}} >}{p_{ICA_{\text{ext}\,n}}} - 1 \,, \quad \text{input to baroreflex: pressure at ICA}_{\text{ext}} \tag{14}$$

$$\alpha_{ECA} = \frac{\alpha_{ECA\,n}}{(1 + \delta d_{ECA})^4} \,, \quad \text{controlled resistance} \tag{15}$$

$$A_{ECA0} = A_{ECA0n} (1 + \delta d_{ECA})^2 \,, \quad \text{controlled area} \tag{16}$$

**Extracranial Arteries/Arterioles ($Ar_{ext}$)**

$$l_{Ar_{\text{ext}}} \frac{\partial A_{Ar_{\text{ext}}}}{\partial t} = f_{Ar_{\text{ext}\,in}} - f_{Ar_{\text{ext}\,out}} \,, \quad \text{continuity} \tag{17}$$

$$p_{ECA} - p_{Ar_{\text{ext}}} = \alpha_{Ar_{\text{ext}}} f_{Ar_{\text{ext}\,in}}\,, \quad \text{momentum} \tag{18}$$

$$p_{Ar_{\text{ext}}} - 0 = E_{Ar_{\text{ext}}} \left( \frac{A_{Ar_{\text{ext}}}}{A_{Ar_{\text{ext}\,0}}} - 1 \right), \quad \text{distensibility} \tag{19}$$

$$\alpha_{Ar_{\text{ext}}} = \frac{\alpha_{Ar_{\text{ext}\,n}}}{(1+\delta d_{ECA})^4}\,, \quad \text{controlled resistance} \tag{20}$$

$$A_{Ar_{\text{ext}\,0}} = A_{Ar_{\text{ext}\,0n}} (1+\delta d_{ECA})^2\,, \quad \text{controlled area} \tag{21}$$

**Extracranial Microcirculation ($\mathbf{Mc_{ext}}$)**

$$l_{Mc_{\text{ext}}} \frac{\partial A_{Mc_{\text{ext}}}}{\partial t} = f_{Mc_{\text{ext}\,in}} - f_{Mc_{\text{ext}\,out}}\,, \quad \text{continuity} \tag{22}$$

$$p_{Ar_{\text{ext}}} - p_{Mc_{\text{ext}}} = \alpha_{Mc_{\text{ext}}} f_{Mc_{\text{ext}\,in}}\,, \quad \text{momentum} \tag{23}$$

$$p_{Mc_{\text{ext}}} - 0 = E_{Mc_{\text{ext}}} \left( \frac{A_{Mc_{\text{ext}}}}{A_{Mc_{\text{ext}\,0}}} - 1 \right), \quad \text{distensibility} \tag{24}$$

**Extracranial Veins ($\mathbf{V_{ext}}$)**

$$l_{V_{\text{ext}}} \frac{\partial A_{V_{\text{ext}}}}{\partial t} = f_{V_{\text{ext}\,in}} - f_{V_{\text{ext}\,out}}\,, \quad \text{continuity} \tag{25}$$

$$p_{Ar_{\text{ext}}} - p_{V_{\text{ext}}} = \alpha_{V_{\text{ext}}} f_{V_{\text{ext}\,in}}\,, \quad \text{momentum} \tag{26}$$

$$p_{V_{\text{ext}}} - 0 = E_{V_{\text{ext}}} \left( \frac{A_{V_{\text{ext}}}}{A_{V_{\text{ext}\,0}}} - 1 \right), \quad \text{distensibility} \tag{27}$$

$$p_{V_{\text{ext}}} - p_{out} = \alpha_{out} f_{V_{\text{ext}\,out}}\,, \quad \text{additional momentum} \tag{28}$$

**Extracranial internal carotid artery ($\mathbf{ICA_{ext}}$)**

$$l_{ICA_{\text{ext}}} \frac{\partial A_{ICA_{\text{ext}}}}{\partial t} = f_{ICA_{\text{ext}\,in}} - f_{ICA_{\text{ext}\,out}}\,, \quad \text{continuity} \tag{29}$$

$$p_{CCA} - p_{ICA_{\text{ext}}} = \alpha_{ICA_{\text{ext}}} f_{ICA_{\text{ext}\,in}}\,, \quad \text{momentum} \tag{30}$$

$$p_{ICA_{\text{ext}}} - 0 = E_{ICA_{\text{ext}}} \left( \frac{A_{ICA_{\text{ext}}}}{A_{ICA_{\text{ext}\,0}}} - 1 \right), \quad \text{distensibility} \tag{31}$$

**Intracranial internal carotid artery ($ICA_{int}$)**

$$l_{ICA_{\text{int}}}\frac{\partial A_{ICA_{\text{int}}}}{\partial t}=f_{ICA_{\text{int}\,in}}-f_{ICA_{\text{int}\,out}}\,,\quad \text{continuity} \tag{32}$$

$$p_{ICA_{\text{ext}}}-p_{ICA_{\text{int}}}=\alpha_{ICA_{\text{int}}}f_{ICA_{\text{int}\,in}}\,,\quad \text{momentum} \tag{33}$$

$$p_{ICA_{\text{int}}}-0.5(p^{R}_{br_{exf}}+p^{L}_{br_{exf}})=E_{ICA_{\text{int}}}\left(\frac{A_{ICA_{\text{int}}}}{A_{ICA_{\text{int}\,0}}}-1\right)\,,\quad \text{distensibility} \tag{34}$$

**Arteries**

$$l^{L,R}_{Ar}\frac{\partial A^{L,R}_{Ar}}{\partial t}=f_{Ar_{in}}^{L,R}-f_{Ar_{out}}^{L,R}\,,\quad \text{continuity} \tag{35}$$

$$p_{ICA}-p^{L,R}_{Ar}=\alpha^{L,R}_{Ar}\,f_{Ar_{in}}^{L,R}\,,\quad \text{momentum} \tag{36}$$

$$p^{L,R}_{Ar}-p^{L,R}_{br_{exf}}=E_{Ar}\left(\frac{A^{L,R}_{Ar}}{A_{Ar_0}^{L,R}}-1\right)\,,\quad \text{distensibility} \tag{37}$$

$$\tau_{Ar}\frac{\partial(\delta d^{L,R}_{Ar})}{\partial t}+\delta d^{L,R}_{Ar}=\sigma_{Ar}(x^{L,R}_{Ar})\,,\quad \text{autoregulation} \tag{38}$$

$$x^{L,R}_{Ar}=\frac{< p^{L,R}_{Ar}-p^{L,R}_{br_{exf}} >}{\left(p^{L,R}_{Ar}-p^{L,R}_{br_{exf}}\right)_n}-1\,,\quad \text{input to autoregulation: transmural pressure} \tag{39}$$

$$\alpha^{L,R}_{Ar}=\frac{\alpha_{Arn}}{(1+\delta d^{L,R}_{Ar})^4}\,,\quad \text{controlled resistance} \tag{40}$$

$$A_{Ar_0}^{L,R}=A_{Ar0n}(1+\delta d^{L,R}_{Ar})^2\,,\quad \text{controlled area} \tag{41}$$

**Arterioles**

$$l^{L,R}_{Al}\frac{\partial A^{L,R}_{Al}}{\partial t}=f_{Al_{in}}^{L,R}-f_{Al_{out}}^{L,R}\,,\quad \text{continuity} \tag{42}$$

$$p^{L,R}_{Ar}-p^{L,R}_{Al}=\alpha^{L,R}_{Al}\,f_{Al_{in}}^{L,R}\,,\quad \text{momentum} \tag{43}$$

$$p^{L,R}_{Al}-p^{L,R}_{br_{exf}}=E_{Al}\left(\frac{A^{L,R}_{Al}}{A_{Al_0}^{L,R}}-1\right)\,,\quad \text{distensibility} \tag{44}$$

$$\tau_{Al}\frac{\partial(\delta d_{Al}^{L,R})}{\partial t}+\delta d_{Al}^{L,R}=\sigma_{Al}(x_{Al}^{L,R})\,,\quad \text{autoregulation} \tag{45}$$

$$x_{Al}^{L,R}=\frac{< p_{Al}^{L,R}-p_{br_{exf}}^{L,R} >}{\left(p_{Al}^{L,R}-p_{br_{exf}}^{L,R}\right)_n}-1\,,\quad \text{input to autoregulation: transmural pressure} \tag{46}$$

$$\alpha_{Al}^{L,R}=\frac{\alpha_{Al\,n}}{(1+\delta d_{Al}^{L,R})^4}\,,\quad \text{controlled resistance} \tag{47}$$

$$A_{Al_0}^{L,R}=A_{Al0n}(1+\delta d_{Al}^{L,R})^2\,,\quad \text{controlled area} \tag{48}$$

**Microcirculation (terminal arterioles, capillaries and venules)**

$$l_{Mc}^{L,R}\frac{\partial A_{Mc}^{L,R}}{\partial t}=f_{Mc_{in}}^{L,R}-f_{Mc_{out}}^{L,R}\,,\quad \text{continuity} \tag{49}$$

$$p_{Al}^{L,R}-p_{Mc}^{L,R}=\alpha_{Mc}^{L,R}f_{Mc_{in}}^{L,R}\,,\quad \text{momentum} \tag{50}$$

$$p_{Mc}^{L,R}-p_{br_{exf}}^{L,R}=E_{Mc}\left(\frac{A_{Mc}^{L,R}}{A_{Mc_0}^{L,R}}-1\right)\,,\quad \text{distensibility} \tag{51}$$

$$\tau_{Mc}\frac{\partial(\delta d_{Mc}^{L,R})}{\partial t}+\delta d_{Mc}^{L,R}=\sigma_{Mc}(x_{Mc}^{L,R})\,,\quad \text{autoregulation} \tag{52}$$

$$x_{Mc}^{L,R}=\frac{< f_{Mc_{in}}^{L,R} >}{\left(f_{Mc_{in}}^{L,R}\right)_n}-1\,,\quad \text{input to autoregulation: blood flow} \tag{53}$$

$$\alpha_{Mc}^{L,R}=\frac{\alpha_{Mc\,n}}{(1+\delta d_{Mc}^{L,R})^4}\,,\quad \text{controlled resistance} \tag{54}$$

$$A_{Mc_0}^{L,R}=A_{Mc0n}(1+\delta d_{Mc}^{L,R})^2\,,\quad \text{controlled area} \tag{55}$$

**Veins**

$$l_{V}^{L,R}\frac{\partial A_{V}^{L,R}}{\partial t}=f_{V_{in}}^{L,R}-f_{V_{out}}^{L,R}\,,\quad \text{continuity} \tag{56}$$

$$p_{Mc}^{L,R}-p_{V}^{L,R}=\alpha_{V}^{L,R}f_{V_{in}}^{L,R}\,,\quad \text{momentum} \tag{57}$$

$$p_{V}^{L,R}-p_{br_{exf}}^{L,R}=E_{V}\left(\frac{A_{V}^{L,R}}{A_{V_0}^{L,R}}-1\right)\,,\quad \text{distensibility} \tag{58}$$

**Venous sinus**

$$l_{vSinus}\frac{\partial A_{vSinus}}{\partial t} = f_{vSinus\,in} - f_{vSinus\,out}\,, \quad \text{continuity} \tag{59}$$

$$0.5(p_V^L + p_V^R) - p_{vSinus} = \alpha_{vSinus} f_{vSinus\,in}\,, \quad \text{momentum} \tag{60}$$

$$p_{vSinus} - p_{out} = \alpha_{out} f_{vSinus\,out}\,, \quad \text{additional momentum} \tag{61}$$

$$p_{vSinus} - 0.5(p^L_{br_{exf}} + p^R_{br_{exf}}) = E_V\left(\frac{A_{vSinus}}{A_{vSinus0}} - 1\right), \quad \text{distensibility} \tag{62}$$

**CSF system**

**Lateral ventricles**

$$l_{Lv}^{L,R}\frac{\partial A_{Lv}^{L,R}}{\partial t} = f_{Lv\,in}^{\;L,R} - f_{Lv\,out}^{\;L,R}\,, \quad \text{continuity} \tag{63}$$

$$p_{Lv}^{L,R} - p_{3V} = \alpha_{3V} f_{Lv\,out}^{\;L,R}\,, \quad \text{momentum} \tag{64}$$

$$p_{Lv}^{L,R} - p_{br_{exf}}^{L,R} = E_{Lv}^{L,R}\left(\frac{A_{Lv}^{L,R}}{A_{Lv_0}^{\;L,R}} - 1\right), \quad \text{distensibility} \tag{65}$$

**Third ventricle**

$$l_{3V}\frac{\partial A_{3V}}{\partial t} = f_{3V\,in} - f_{3V\,out}\,, \quad \text{continuity} \tag{66}$$

$$p_{3V} - 0.5(p^L_{br_{exf}} + p^R_{br_{exf}}) = E_{3V}\left(\frac{A_{3V}}{A_{3V\,0}} - 1\right), \quad \text{distensibility} \tag{67}$$

**Fourth ventricle**

$$l_{4V}\frac{\partial A_{4V}}{\partial t} = f_{4V\,in} - f_{4V\,out}\,, \quad \text{continuity} \tag{68}$$

$$p_{3V} - p_{4V} = \alpha_{4V} f_{4V\,in}\,, \quad \text{momentum} \tag{69}$$

$$p_{4V} - 0.5(p^L_{br_{exf}} + p^R_{br_{exf}}) = E_{4V}\left(\frac{A_{4V}}{A_{4V\,0}} - 1\right), \quad \text{distensibility} \tag{70}$$

**Cranial subarachnoid space**

$$l_{cSAS}\frac{\partial A_{cSAS}}{\partial t} = f_{cSAS\,in} - f_{cSAS\,out}\,, \quad \text{continuity} \tag{71}$$

$$p_{4V} - p_{cSAS} = \alpha_{cSAS} f_{cSAS\,in}\,, \quad \text{momentum} \tag{72}$$

$$p_{cSAS} - 0.5(p^{L}_{br_{exf}} + p^{R}_{br_{exf}}) = E_{cSAS}\left(\frac{A_{cSAS}}{A_{cSAS0}} - 1\right)\,, \quad \text{distensibility} \tag{73}$$

**Spinal subarachnoid space**

$$l_{sp.canal}\frac{\partial A_{sp.canal}}{\partial t} = f_{sp.canal\,in} - f_{sp.canal\,out}\,, \quad \text{continuity} \tag{74}$$

$$p_{cSAS} - p_{sp.canal} = \alpha_{sp.canal} f_{sp.canal\,in}\,, \quad \text{momentum} \tag{75}$$

$$p_{sp.canal} - 0 = E_{sp.canal}\left(\frac{A_{sp.canal}}{A_{sp.canal\,0}} - 1\right)\,, \quad \text{distensibility} \tag{76}$$

**Brain parenchyma**

$$f_{br_{exf}\,in}^{\;L,R} - f_{br_{exf}\,out}^{\;L,R} = 0\,, \quad \text{continuity} \tag{77}$$

$$f_{br_{exf}\,in}^{\;L,R} = S^{L,R}_{const_{Mc\to br}} + S^{L,R}_{Mc\to br} \tag{78}$$

$$f_{br_{exf}\,out}^{\;L,R} = S^{L,R}_{const_{br\to Lv}} + S^{L,R}_{br\to Lv} \tag{79}$$

$$p^{L,R}_{Mc} - p^{L,R}_{br_{exf}} = \alpha_{Mc\to br} S^{L,R}_{Mc\to br}\,, \quad \text{momentum} \tag{80}$$

$$p^{L,R}_{br_{exf}} - p^{L,R}_{Lv} = \alpha_{br\to Lv} S^{L,R}_{br\to Lv}\,, \quad \text{momentum} \tag{81}$$

$$S^{L,R}_{const_{Mc\to br}} = 0.0005 \text{ ml/s} \tag{82}$$

$$S^{L,R}_{const_{br\to Lv}} = 0.0005 \text{ ml/s} \tag{83}$$

Monro-Kellie doctrine is enforced separetely at each side of the skull:

$$\begin{aligned} &V^{L,R}_{total_{intracranial}} = \text{constant} \Rightarrow \\ &(0.5V_{ICA_{\text{int}}} + V^{L,R}_{Ar} + V^{L,R}_{Al} + V^{L,R}_{Mc} + V^{L,R}_{V} + V_{vSinus}) \\ &\qquad + (V^{L,R}_{Lv} + 0.5V_{3V} + 0.5V_{4V} + 0.5V_{cSAS}) + (V^{L,R}_{br_{solid}} + V^{L,R}_{br_{exf}}) = \text{constant} \end{aligned} \tag{84}$$

Equations connecting the compartments and source terms

$$f_{CCA\,out} = f_{ICA_{\text{ext}}\,in} + f_{ECA\,in} \tag{85}$$

$$f_{ECA\,out} = f_{Ar_{\text{ext}}\,in} \tag{86}$$

$$f_{Ar_{\text{ext}}\,out} = f_{Mc_{\text{ext}}\,in} \tag{87}$$

$$f_{Mc_{\text{ext}}\,out} = f_{V_{\text{ext}}\,in} \tag{88}$$

$$f_{ICA_{\text{ext}}\,out} = f_{ICA_{\text{int}}\,in} \tag{89}$$

$$f_{ICA_{\text{int}}\,out} = f_{Ar}{}_{in}^{L} + f_{Ar}{}_{in}^{R} \tag{90}$$

$$f_{Ar}{}_{out}^{L,R} = f_{Al}{}_{in}^{L,R} \tag{91}$$

$$f_{Al}{}_{out}^{L,R} = f_{Mc}{}_{in}^{L,R} \tag{92}$$

$$f_{Mc}{}_{out}^{L,R} = f_{Lv}{}_{in}^{L,R} + f_{br_{exf}}{}_{in}^{L,R} + f_{V}{}_{in}^{L,R} \tag{93}$$

$$f_{vSinus\,in} = f_{V}{}_{out}^{L} + f_{V}{}_{out}^{R} + f_{reabsorption} \tag{94}$$

$$f_{Lv}{}_{in}^{L,R} = S_{const_{Mc\to Lv}}^{L,R} \tag{95}$$

$$S_{const_{Mc\to Lv}}^{L,R} = 0.003 \text{ ml/s} \tag{96}$$

$$f_{3V\,in} - f_{Lv}{}_{out}^{L} - f_{Lv}{}_{out}^{R} = f_{br_{exf}}{}_{out}^{L} + f_{br_{exf}}{}_{out}^{R} \tag{97}$$

$$f_{4V\,in} = f_{3V\,out} \tag{98}$$

$$f_{4V\,out} = f_{cSAS\,in} \tag{99}$$

$$f_{cSAS_{out}} = f_{sp.canal_{in}} + f_{reabsorption} \tag{100}$$

$$f_{reabsorption} = k(p_{cSAS} - p_{vSinus}) \tag{101}$$

$$1/k = 15.62 \text{ mmHg·min/ml} \tag{102}$$

$$f_{sp.canal_{out}} = 0 \tag{103}$$

$$p_{out} = 2.5 \text{ mmHg, zero amplitude} \tag{104}$$

$$\alpha_{out} = 0.0809088 \text{ mmHg·s/ml} \tag{105}$$

#### A2. Parameters used in the model.

| Compartment | Length (cm) | Area at rest (cm²) | Elastance (mmHg) | Resistance (mmHg·s/ml) |
|---|---|---|---|---|
| Common carotid | (12.14 + 8.13)/2* | 0.5765* | 1600 | 0.04854525* |
| External carotid | 6.10* | 0.3223* | 800000* | 0.1220* |
| Extracranial arteries | 6.16* | 12.29* | 1920 | 6.433* |
| Extracranial microcirculation | 0.2730* | 122.6* | 6800 | 13.638* |
| Extracranial veins | 7.473* | 17.23* | 756.56 | 3.1104* |
| Extracranial internal carotid | 8.6* | 0.3947* | 800 | 0.0786* |
| Intracranial internal carotid | 9.1* | 0.1973* | 1600 | 0.0831* |
| Arteries | 4.15 | 3.42 | 1920 | 2.76912 |
| Arterioles | 1.75 | 4.74 | 410.28 | 4.33831545 |
| Microcirculation | 0.2618 | 38.0 | 6800 | 6.002803222 |
| Veins | 7.165 | 5.3388 | 756.56 | 2.078301471 |
| Venous sinus | 15.0 | 0.86 | 90.007 | 0.161896 |
| Lateral ventricles | 0.75 | 12.0 | 10.0 | 500.0 ($\alpha_{br \to Lv}$) |
| Third ventricle | 1.0 | 2.5 | 10.0 | 1.0 |
| Fourth ventricle | 1.0 | 3.5 | 10.0 | 1.0 |
| Cranial subarachnoid space | 1.69 | 17.7658 | 80.0 | 1.0 |
| Spinal cord | 43.0 | 2.0 | 300.0 | 0.1 |
| Brain extracellular fluid | 7.0 | 30.0 | ——— | 8152.42 ($\alpha_{Mc \to br}$) |
| Brain solid cell matrix | 7.0 | 70.0 | ——— | ——— |

**Table S5.** Baseline values used in the model for each compartment. All parameters with the exception of * were taken from published and unpublished data[35]

.

| Parameter | Hypotension | Hypertension |
|---|---|---|
| CBF = $f_{ICA_n}$ | 12.518 ml/s | 12.515 ml/s |
| $(p_{ICA_{ext}})_n$ | 100.591 mmHg | 100.599 mmHg |
| $\left(p^{L,R}_{Ar} - p^{L,R}_{br_{exf}}\right)_n$ | 75.085 mmHg | 70.464 mmHg |
| $\left(p^{L,R}_{Al} - p^{L,R}_{br_{exf}}\right)_n$ | 47.935 mmHg | 43.317 mmHg |
| $\left(f^{L,R}_{Mc_{in}}\right)_n$ | 6.256 ml/s | 6.257 ml/s |
| $HR_n$ | 1 Hz = 60 bpm | 1 Hz = 60 bpm |
| $p^{L,R}_{br_{exf}}$ | 7.133 mmHg | 11.766 mmHg |

**Table S6.** Baseline values for pressures, flows and HR used at hypotension and hypertension simulations.

| Age (years) | Pressure wave Fourier coefficients [mmHg] |
|---|---|
| 19 | $(a_0, ..., a_8)$ = (+72.805, -1.7232, -0.2418, -3.7949, -0.9176, -1.0510, -0.9843, -0.3240, -0.4260)<br>$(b_1, ..., b_8)$ = (+7.0992, +5.0640, +0.9592, -0.2273, +0.4517, -0.3920, -0.2460, -0.1824) |
| 42 | $(a_0, ..., a_8)$ = (+98.529, -4.2922, -4.9021, -2.4110, -0.8043, -0.7461, -1.3538, -0.8557, -0.3375)<br>$(b_1, ..., b_8)$ = (+16.0276, +3.586, +0.5741, +0.3451, +0.8614, +0.3377, -0.4699, -0.3575) |
| 83 | $(a_0, ..., a_8)$ = (+99.341, -8.0572, -12.1456, -3.1937, -0.7437, -1.2039, -1.2926, -0.5074, -0.5952)<br>$(b_1, ..., b_8)$ = (+ 31.713, +3.1270, -1.6578, -0.0335, + 0.6687, -0.2084, -0.3016, -0.1075) |

**Table S7.** Fourier coefficients for aortic pressure waves fitted for individuals aging 19, 42 and 83 years[67].
$p_{init} = a_0 + \sum_{n=1}^{8} a_n cos(n\omega x) + b_n sin(n\omega x)$

##### A3. Simulation setup.

The model was implemented in Julia[68] v.1.5.2 as a differential-algebraic system of 120 equations. The derivatives were calculated using a first-order backward difference approximation and the system was solved step-by-step, the solution of the previous time step being the starting point for the next one. The time step was 0.01 s for the simulations of acute hypotension, random input and to study the effects of the arterial pressure wave, 0.05 s for hypertension and 0.1 s for the static autoregulation analysis. We used the nonlinear solver nlsolve() from the NLsolve.jl package.
